# A Graph-Directed Approach for Creation of a Homology Modeling Library: Application to Venom Structure Prediction

**DOI:** 10.1101/828129

**Authors:** Rachael A. Mansbach, Srirupa Chakraborty, Timothy Travers, S. Gnanakaran

## Abstract

Many toxins are short, cysteine-rich peptides that are of great interest as novel therapeutic leads and of great concern as lethal biological agents due to their high affinity and specificity for various receptors involved in neuromuscular transmission. To perform initial candidate identification for design of a drug impacting a particular receptor or for threat assessment as a harmful toxin, one requires a set of candidate structures of reasonable accuracy with potential for interaction with the target receptor. In this article, we introduce a graph-based algorithm for identifying good extant template structures from a library of evolutionarily-related cysteine-containing sequences for structural determination of target sequences by homology modeling. We employ this approach to study the conotoxins, a set of toxin peptides produced by the family of aquatic cone snails. Currently, of the approximately six thousand known conotoxin sequences, only about three percent have experimentally characterized three-dimensional structures, leading to a serious bottleneck in identifying potential drug candidates. We demonstrate that the conotoxin template library generated by our approach may be employed to perform homology modeling and greatly increase the number of characterized conotoxin structures. We also show how our approach can guide experimental design by identifying and ranking sequences for structural characterization in a similar manner. Overall, we present and validate an approach for venom structure modeling and employ it to expand the library of extant conotoxin structures by almost 300% through homology modeling employing the template library determined in our approach.

## 1 Introduction

Toxins have for a long time been considered a rich natural source of therapeutic leads because of their high specificity and binding affinity for various receptors involved in different biological pathways [Zambelli et al., 2016, Verdes et al., 2016]. The drug ziconotide, for example, is a potent analgesic derived from a toxin produced by the aquatic cone snail species *Conus magus* [Miljanich, 2004]. The on-average smaller size of toxins–typically < 100 amino acids along with a sizeable proportion < 30 amino acids long [Dang, 2019]–means they can be employed with relative ease in high-throughput *in silico* screenings to rationally identify candidates for initial scaffolds interacting with a particular receptor of interest. Indeed, this is a fruitful initial line of inquiry for improving drug discovery outcomes and productivity [Romano and Tatonetti, 2019]: in one recent study of note the authors employed a docking approach to identify *α*-conotoxin BuIA, produced by species *Conus bullatus*, as a competitive agonist for the lysophosphatidic acid receptor 6, a G-protein coupled receptor involved in the development of several cancers [Younis and Rashid, 2017]. However, such an approach is limited by the necessity of possessing a library of at least moderately-accurate structures of potential toxin candidates [Sledz and Caflisch, 2018]: more structures mean a larger search space and hence a higher likelihood of identifying good initial leads.

Aside from their therapeutic benefit, toxins such as these also pose a threat to biosecurity. The high-throughput evaluation of toxin mode of action as well as the diagnosis and decontamination of disulfide rich toxins, either natural or man-made, is required for public health safety. Rapid advances in synthetic biology have created challenges in determining the health risks posed by natural toxins or modified toxins with even higher pathogenicity [Gomez-Tatay and Hernandez-Andreu, 2019]. Thus, in tandem with high-throughput screening for therapeutic design, it is necessary to also be able to perform high-throughput screening for threat identification and determination of toxin targets and mechanisms of action. Structural characterization stands as a rate-limiting step for high-throughput screening for both therapeutic design and toxin threat characterization, as identified sequences often far outnumber determined structures. For example, only about 3% of sequences isolated from cone snail venom have corresponding experimentally-determined structures [Mansbach et al., 2019].

If the structures of proteins could be rapidly predicted strictly from their sequences, structural determination would not be a bottleneck; however, structure prediction from sequence still remains a challenging proposition [Huang et al., 2016]. Ab initio or de novo modeling approaches for obtaining protein structure predictions by modeling essential folding physics are prohibitively expensive except for small proteins of about 20-62 residues in size [Pitera and Swope, 2003, Ensign et al., 2007, Voelz et al., 2010, Sborgi et al., 2015]. Even for proteins short enough to be de novo modelled in isolation, this can become expensive if a large number of different structures are desired. Structure prediction for a query sequence becomes more tractable when experimentally-resolved structures are available for evolutionarily related sequences: this is referred to as homology modeling [Dill and MacCallum, 2012]. For typical proteins (at least 100 amino acids long), a useful rule of thumb for building a homology model of a protein with unknown structure using a structurally characterized protein as the template is that both proteins should share at least 25% sequence identity [Baker and Sali, 2001, Xiang, 2006]. Of note, several protein structure prediction algorithms use a combination of ab initio and homology modeling approaches. For instance, both I-TASSER [Yang et al., 2015, Zhang, 2008] and ROSETTA [Bradley et al., 2005, Simons et al., 1997] split up the query sequence into fragments that will be searched against a library to identify structure fragments (the homology modeling part), followed by assembly of these fragments into a full-length structural model via molecular dynamics or Monte Carlo simulations using either physics-based or knowledge-based force fields (the ab initio modeling part).

One challenge in applying homology modeling to toxins is that, due to their on-average smaller size, the so-called modeling “safe zone” where structural similarity can be inferred requires a higher sequence identity during structural template selection than the 25% rule of thumb for typical proteins [Krieger et al., 2003, Kong et al., 2004]. This is because at shorter peptide lengths, a sequence identity of 25% is more likely to have arisen by random chance and not due to any evolutionary constraints on the structure. A related challenge occurs in constructing suitable template libraries that contain sufficient information for building accurate homology models for these short peptides. To apply the homology modeling framework for shorter peptides, a reasonable heuristic instead becomes that the alignment length and percent identity fall above the phenomenological curve introduced by Rost [Rost, 1999] (see Eqn. 1 and Fig. S1). The relative steepness of the Rost curve for alignment lengths of less than fifty amino acids provides an illustration of why, for peptides of such lengths, it is important to use the actual functional form, rather than a static cutoff, to assess whether a pairwise alignment contains sufficient information for homology modeling.

In this article, we propose the use of a simple graph-based algorithm for homology modeling of toxins. Graph theory has a long and storied history of usage for sequence-grouping tasks such as homology detection [Santiago et al., 2018], structure prediction [Bolten et al., 2001, Pipenbacher et al., 2002, Yan et al., 2011], protein family identification [Abascal and Valencia, 2002, Enright and Ouzounis, 2000], and even direct homology modeling [Yan et al., 2011]. For large heterogeneous databases, it can be challenging to identify homologues and a number of sophisticated algorithms have been developed for such purposes; we instead focus on the problem of homology modeling a set of cysteine-rich toxins known to be evolutionarily related. In our approach, we employ the number and placement of cysteines within a sequence as a rough initial estimate of functional and structural relatedness. We apply our approach to the so-called conotoxins, which are small, cysteine-rich peptide toxins produced by the cone snails [Uribe et al., 2018, Mansbach et al., 2019].

In the following sections, we present our graph-based approach and employ it to construct sequence graphs and identify good libraries of templates for homology modeling. We demonstrate that these libraries improve outcomes for structure homology modeling over using the typical 25% flat cutoff and employ them as part of a homology modeling procedure that results in a significantly expanded library of structures for the conotoxins that will be of use in future high-throughput studies. In addition, we use the graph-based approach to construct a set of tables indicating sequences whose experimental or ab initio structural characterization is predicted to be most valuable in creating a broad structure library by using homology modeling.

## 2 Results

We initialize the algorithm by separating a set of over 2000 known conotoxin sequences into databases containing four, six, eight, and ten cysteines respectively. For each database, we construct graphs of sequences in which an edge between two nodes (i.e., sequences) represents a pairwise alignment that is of sufficient length and percent identity to fall into the safe homology modeling zone above the Rost curve (cf. Eqn. 1 and Fig. S1). Some portion of the sequences have known structures, such that the corresponding nodes are annotated with the relevant PDB ID(s). We employ the graphs thus generated to iteratively add nodes with structures to a library of templates for homology modeling (see Fig. 1 for a schematic illustration of the procedure). We term this set of sequences {ℒ_ex_}, the set of existing structural library templates (cf. Fig. 2). Nodes are added to {ℒ_ex_} in a greedy manner, in order of highest node degree, such that the resulting library will contain enough templates to homology model as many non-structurally-characterized sequences as possible but with small sequence overlap and retaining a number of non-library structures for quality assessment. Since this is approximately the vertex-covering problem of a graph, we cannot find a globally optimal solution, as that problem is NP-complete [Karp, 1972]. We halt the procedure once either we have no further nodes with structures to add or there are no remaining sequences in a given connected component of the graph that are not connected to at least one library template sequence, such that all sequences in that component may be structurally characterized by homology modeling. We refer to the set of sequences that may be homology modeled based on set {ℒ_ex_} as set {𝒞 (ℒ_ex_}) that are covered by {ℒ_ex_}. We also perform a similar procedure–but without the constraint of structure annotation–on the nodes absent {ℒ_ex_} to identify the set {ℒ_proj_} that are of interest for experimental or ab initio structural characterization such that they cover the remaining set {𝒞 (ℒ_proj_}).

**Figure 1:**
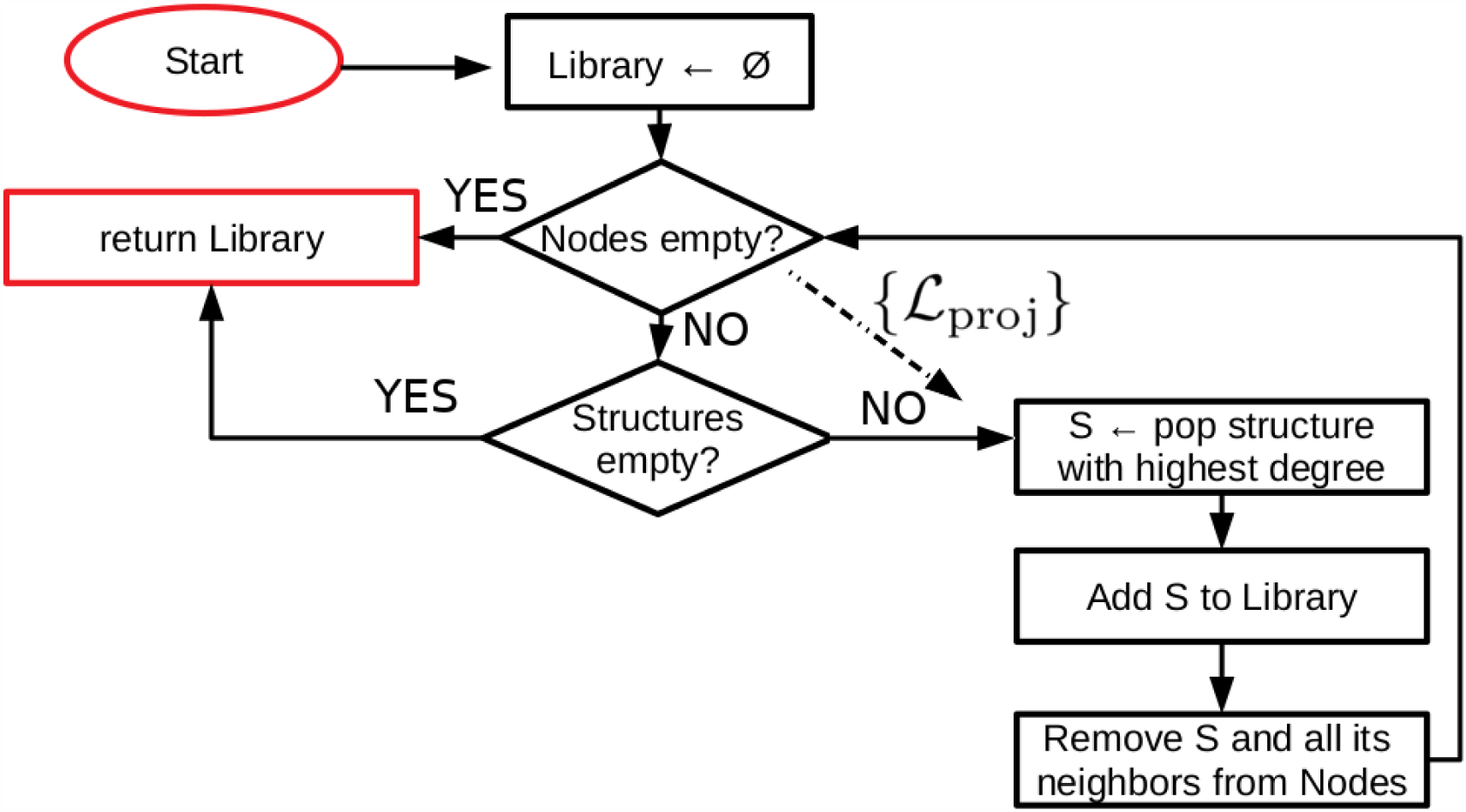
Schematic of a simple graph-based algorithm for constructing a library of structural templates for homology modeling. For each connected component in the graph of sequences, where an edge represents the ability to homology model one sequence based on another, we employ a greedy approach to find a good library of template structures that cover as much of the sequence space as possible. For computation of the sequence set {ℒ_proj_} of interest for experimental or ab initio characterization, we skip consideration of the structures and run the algorithm on the subset with structure-associated sequences removed.

**Figure 2:**
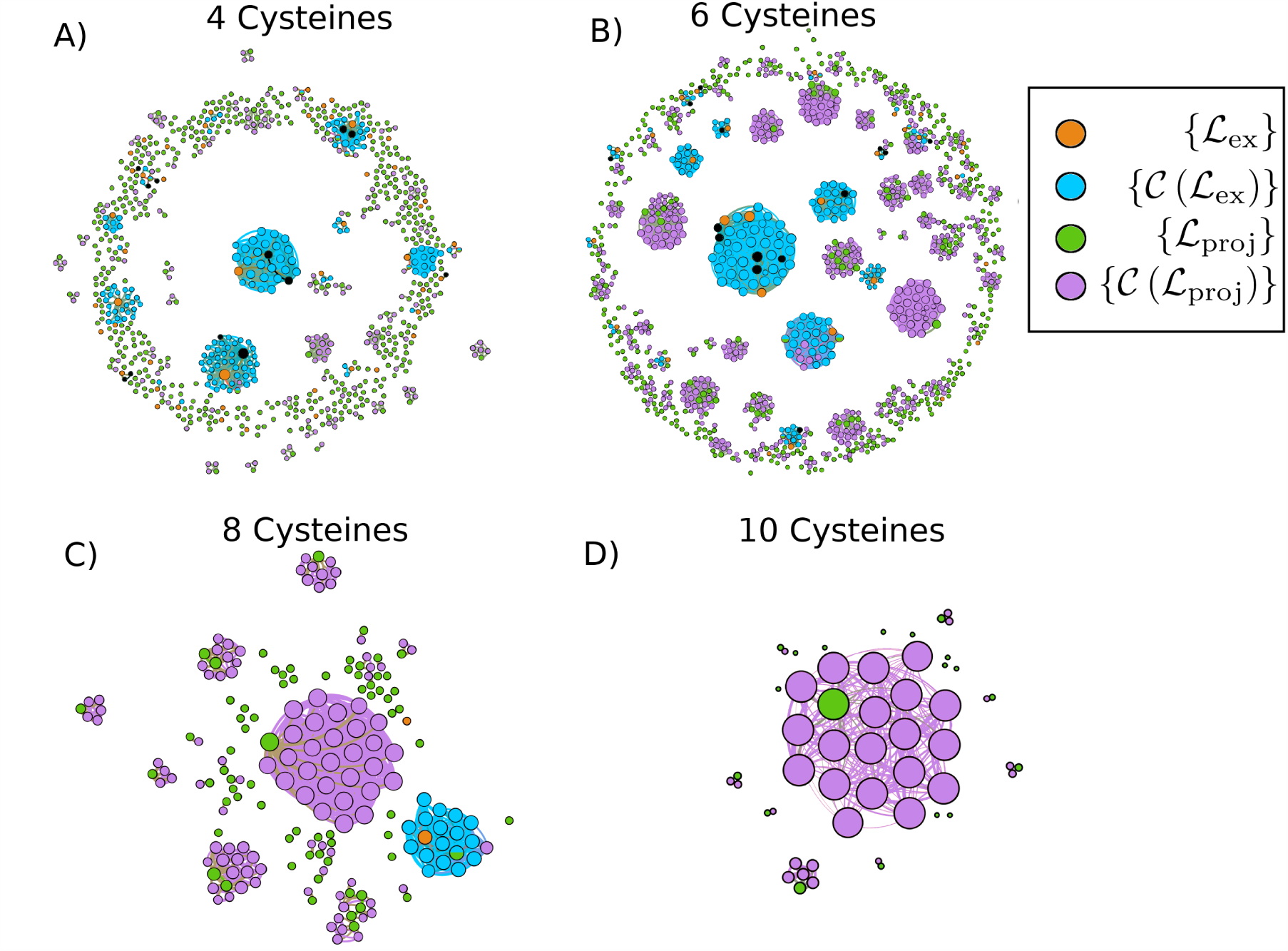
Graph of conotoxins containing (A) four cysteines, (B) six cysteines, (C) eight cysteines and (D) ten cysteines where nodes are sequences and edges exist between sequences with pairwise alignments that have high enough length and percent identity to fall above the Rost curve with *n* = 5% (Eqn. 1). We show the set {ℒ_ex_} of sequences added to the template libraries in orange, the set of sequences corresponding to unselected structures in black, the set of covered sequences {𝒞 (ℒ_ex_}) that we homology model based on the templates included in the library in blue, and the set of projected sequences {ℒ_proj_} in green whose structures are in need of characterization in order that the rest of the sequences {𝒞 (ℒ_proj_}) in magenta may be homology modeled based on some template. The sizes of the nodes corresponds to their degree; that is the number of other sequences that they can be modeled based on or used to model. Node locations and edge lengths were chosen for ease of visualization of separate connectec components. Visualization of the graphs was produced with Gephi 0.9.2 [Bastian et al., 2009].

In Fig. 2, we present the sequence graphs for sets of conotoxin sequences with four, six, eight, and ten cysteines respectively. We specifically display the set {ℒ_ex_} (in orange), which we employ to predict structures for the set {𝒞 (ℒ_ex_}) (in blue) by homology modeling. We show {ℒ_proj_} (in green) whose structural characterization from either experiment or ab initio modeling would lead to coverage by homology modeling of the set {𝒞 (ℒ_proj_}) (in magenta) that comprises sequences with no characterized structure and not covered by set {ℒ_ex_}.

These figures demonstrate that we are able to characterize a large number (and moderate proportion) of unknown conotoxin structures, which may be used for high throughput screening. Specifically, out of the 801 sequences with four cysteines, 61 (7.6% of total) currently have experimentally-resolved structures. The graph-based approach selected 49 (6.1% of total) of these structures as comprising the four cysteine template library (set {ℒ_ex_}; orange circles in Fig. 2a, while the unselected structures are represented in black), which allowed for homology modeling of a further 143 (17.9% of total) sequences (set {𝒞 (ℒ_ex_}); blue circles in Fig. 2a). This corresponds to an increase of over 230% for the number of structurally characterized sequences over the original 61. In addition, the graph-based approach indicated a further 453 sequences (56.6% of total) would need to be characterized, experimentally or ab initio, to allow for homology modeling of the remaining 151 (18.9% of total). Out of the 1,113 sequences with six cysteines, 44 (4.0% of total) currently have experimentally-resolved structures. The graph-based approach selected 30 (2.7% of total) of these structures as comprising the six cysteine template library, which allowed for homology modeling of a further 148 (13.3% of total) sequences. This corresponds to an increase of over 330% for the number of structurally characterized sequences over the original 44. In addition, the graph-based approach indicated a further 419 sequences (37.6% of total) would need to be characterized, experimentally or ab initio, to allow for homology modeling of the remaining 509 (45.7% of total). Out of the 190 sequences with eight cysteines, 2 (1.1% of total) currently have experimentally-resolved structures and were selected as comprising the entire template library, which allowed for homology modeling of a further 17 (8.9% of total) sequences. This corresponds to an increase of 850% for the number of structurally characterized sequences over the original 2. In addition, the graph-based approach indicated a further 71 (37.4% of total) would need to be characterized, experimentally or ab initio, to allow for homology modeling of the remaining 101 (53.1% of total). There are no known structures corresponding to ten cysteine sequences so there is no current coverage. The graph-based approach indicated that 19 of the total 53 sequences (35.8%) would have to be characterized to allow for homology modeling of the remaining 34 (64.2%). In Tables 1, 2, and 3, we list all PDB IDS of the structures included in the template set {ℒ_ex_} constructed for the four and six cysteine sequences, along with their sequences and the name or names of the corresponding toxins.

**Table 1:**
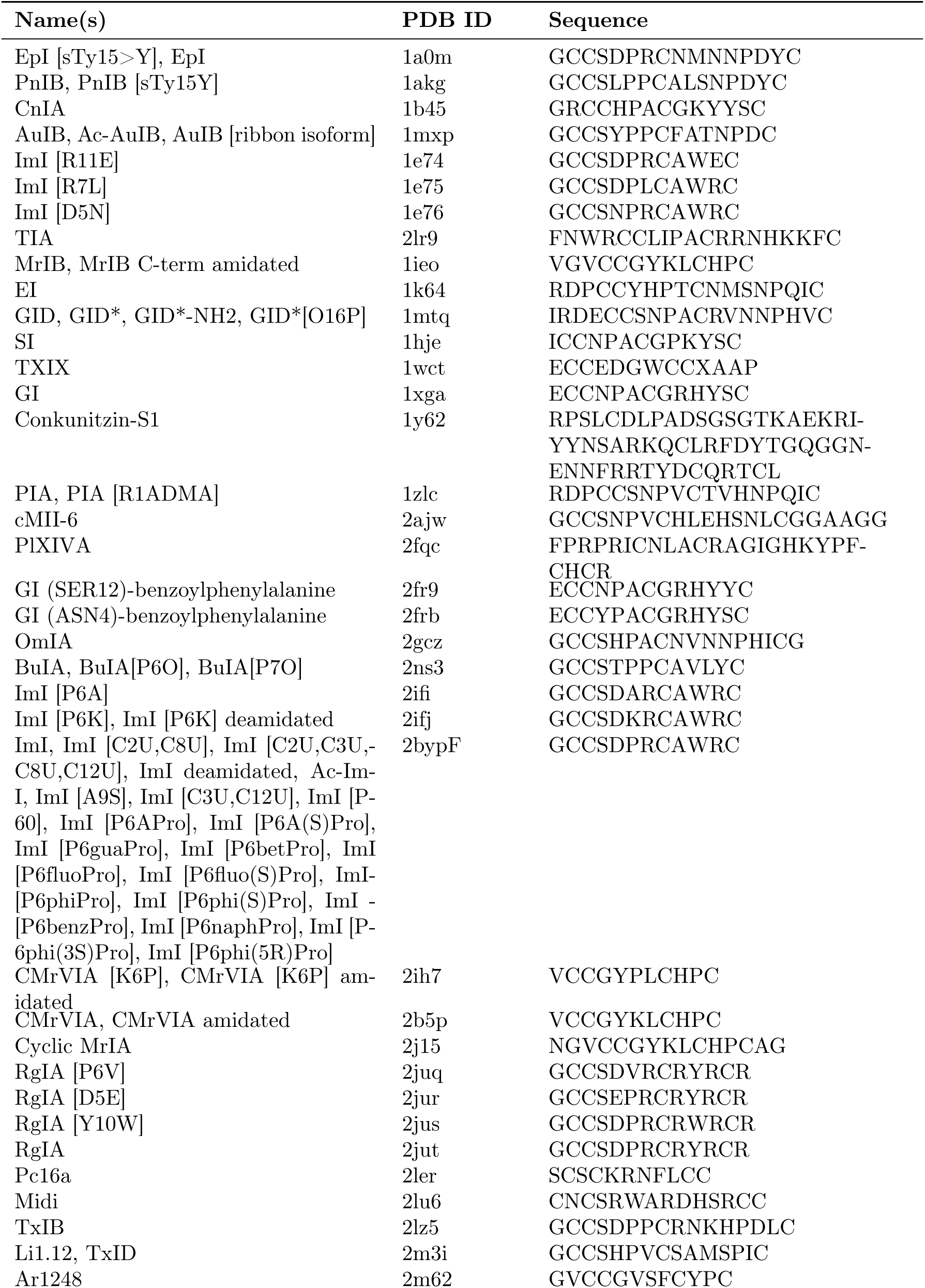
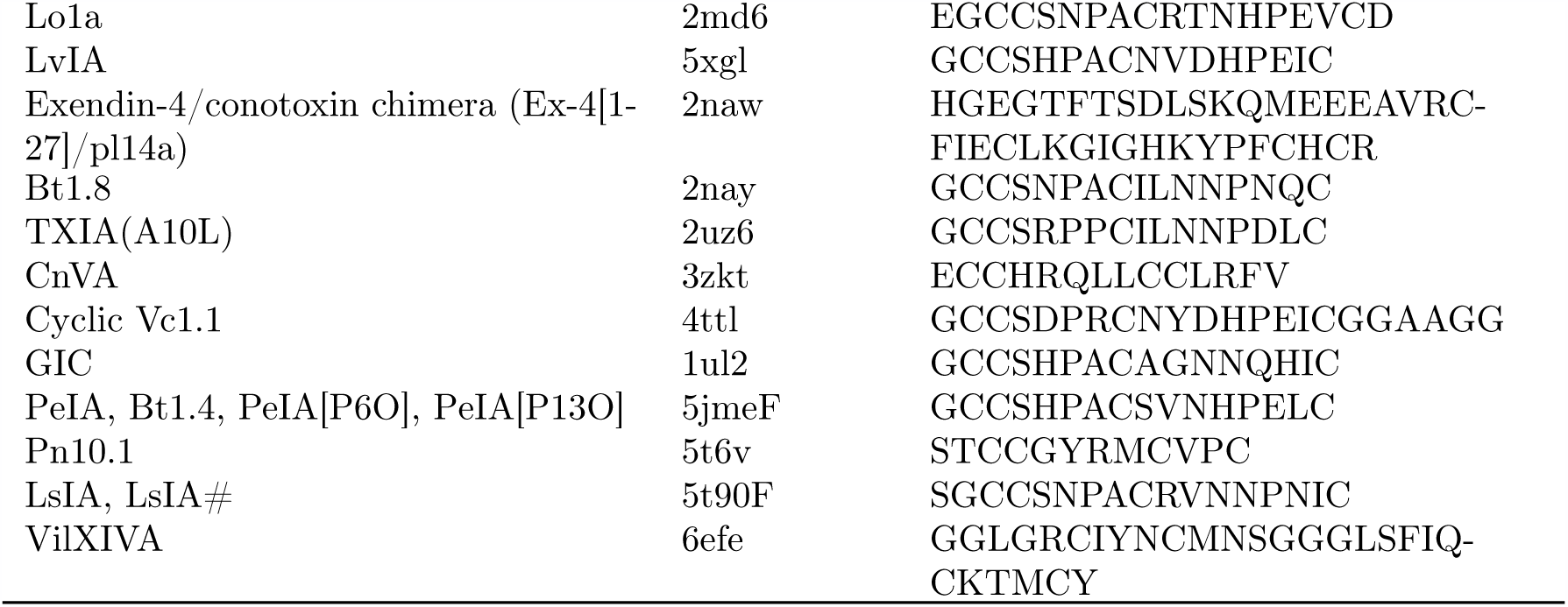
List of conotoxins with corresponding PDB structure IDs [Berman et al., 2000] comprising 4C library. Name or names of sequences are taken from the Conoserver database [Kaas et al., 2012]. Multiple names for the same sequence indicate the same sequence is produced by different species or has different post-translational modifications.

**Table 2:**
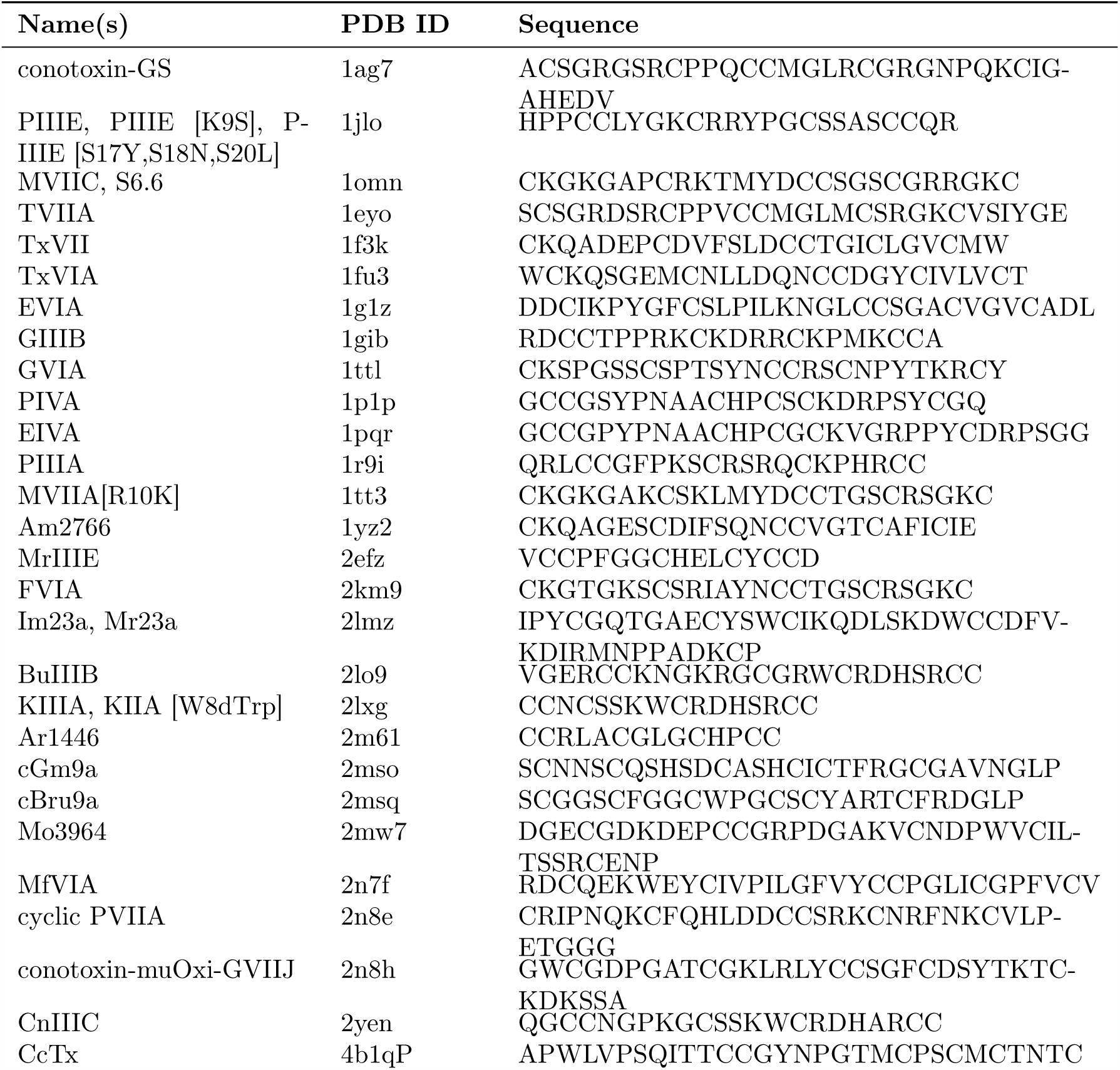
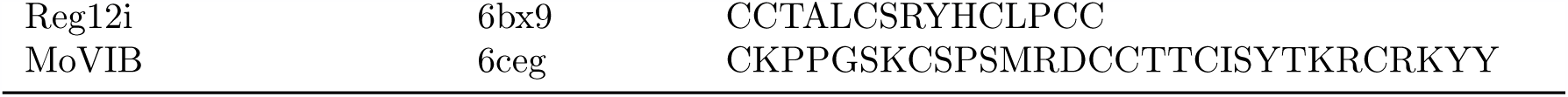
List of conotoxins with corresponding PDB structure IDS [Berman et al., 2000] comprising 6C library. Name or names of sequences are taken from the Conoserver database [Kaas et al., 2012]. Multiple names for the same sequence indicate the same sequence is produced by different species or has different post-translational modifications.

**Table 3:** List of conotoxins with corresponding PDB structure IDS [Berman et al., 2000] comprising 8C library. Name or names of sequences are taken from the Conoserver database [Kaas et al., 2012]. Multiple names for the same sequence indicate the same sequence is produced by different species or has different post-translational modifications.

| Name(s) | PDB ID | Sequence |
| --- | --- | --- |
| G11.1 | 6cei | CAVTHEKCSDDYDCCGSLCCVGICAKTIAPCK |
| RXIA, RXIA [Btr33>W] | 2p4l | GPSFCKADEKPCEYHADCCNCCLSGICAPSTN-<br>WILPGCSTSSFFKI |

In Fig. S2, we present the same sequence graphs used to construct the template libraries, but we color the nodes by relative sequence length instead of set occupation. A significant proportion of isolated sequences (nodes with no connections that therefore cannot be homology modeled) are relatively short (cf. the ring of small red nodes in Fig. S2A and to a lesser extent in Fig. S2B), which demonstrates that a high proportion of isolated nodes may be characterized well through rapid ab initio modeling, particularly for the four and six cysteine sequences. Specifically, 372 of the 453 four cysteine {ℒ_proj_} sequences (82.1%) are isolated nodes with no edges; of these 298 (80.1%) are shorter than 20 amino acids, and 353 (94.9%) are shorter than 30 amino acids in length. In addition, 239 of the 419 six cysteine {ℒ_proj_} sequences (57.0%) are isolated nodes; of these 86 (36.0%) are shorter than 20 amino acids, and 163 (68.2%) are shorter than 30 amino acids in length. Conversely, 41 of the 71 eight cysteine {ℒ_proj_} sequences (57.7%) are isolated nodes, and of these only 5 (12.2%) are shorter than 30 amino acids in length; 10 of the 19 ten cysteine {ℒ_proj_} sequences (52.6%) are isolated nodes and of these none are shorter than 30 amino acids in length.

In Fig. 3, we assess the quality of the template libraries for homology modeling, constructed using the graph-based approach employing the Rost cutoff, and compared with a set of template libraries based on a static 25% rule-of-thumb cutoff. In Fig. 3A-B, we constructed homology models for each structure in a library using the other structures in that library and computed the root-mean-square deviation (RMSD) between each modeled structure with the corresponding experimental structure. In Fig. 3C-D, a similar assessment was performed for all structures that were not included in each template library. As expected, there is a statistically significant improvement (downwards shift in the distribution, two-tailed Kolmogorov-Smirnov test with *p <* 0.05) for both in- and out-of-library structures when using the Rost cutoff as compared to the 25% cutoff, which verifies the necessity of our approach. For in-library assessment, the mean of the distribution drops from 4.0 ± 0.7 Å to 1.5 ± 0.2 Å for the four cysteine library and from 3.8 ± 0.6 Å to 2.1 ± 0.2 Å for the six cysteine library, while for out-of-library assessment, the mean of the distribution drops from 1.7 ± 0.1 Å to 1.0 ± 0.2 Å for the four cysteine library and from 1.82 ± 0.09 Å to 1.4 ± 0.1 Å for the six cysteine library.

**Figure 3:**
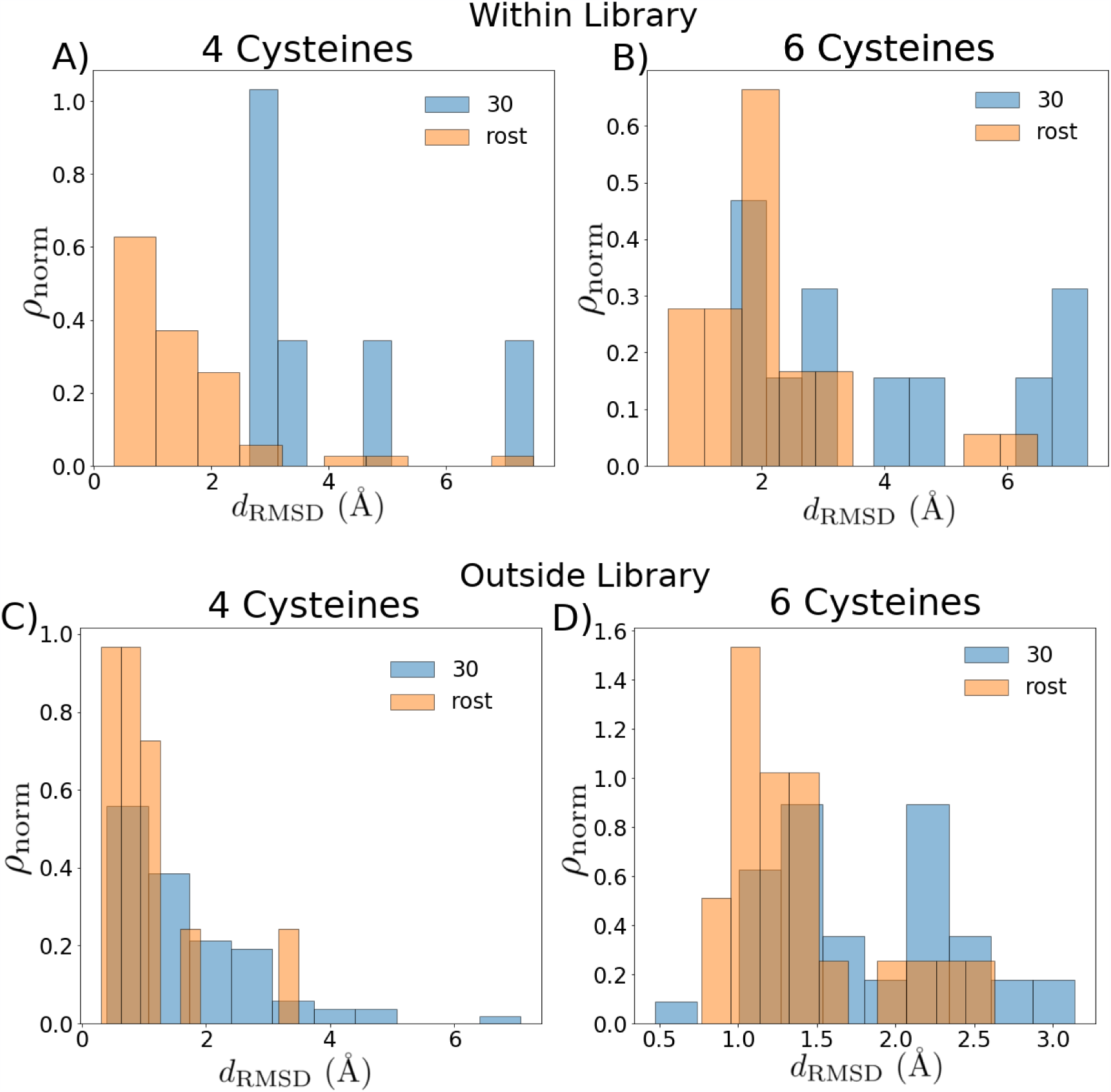
Quality of graph-based template library selection criteria. Comparison of root-mean-square deviation (RMSD) distributions from experimental structures for (A-B) structures within the libraries, with each structure modeled by selecting from all other templates within the given library, and (C-D) structures outside the libraries modeled by selecting from all templates within the given library. For each homology modeled structure, we choose the best fit to experiment. The distributions produced by the simple 30% cutoff libraries are shown in blue; the distributions produced by using the graph-based algorithm are shown in orange.

To illustrate the reason that a static cutoff is less accurate, in Fig. S3, we display an approximation of the distribution of minimum percent identity needed to construct a reliable homology model for a given conotoxin sequence. The minimum required percent identity varies greatly for different conotoxins: although almost none of the conotoxins are long enough to employ the typical 25% cutoff, the relatively large width of the distributions even among conotoxins with the same number of cysteines indicates that choosing a static cutoff is not appropriate. A static cutoff could impact modeling accuracy by either underestimating the needed percent identity for a short sequence or by overestimating the needed percent identity for a long one and thus removing from consideration templates that would otherwise be appropriate, although in the case of a set of short sequences like the conotoxins the primary source of loss of accuracy is expected to be the former.

In Tables 4 and 5 and Tables S1 and S2, we present and rank the set of conotoxins that are of greatest interest for experimental characterization, as availability of experimental structures for these sequences (belonging to set {ℒ_proj_}) would allow homology modeling of the remainder of the (nonisolated) sequences (belonging to set {𝒞 (ℒ_proj_})). We present these in order of greatest graph degree, since greater degree in the graph corresponds to the ability to cover a greater number of sequences. Thus, we suggest that experimental structural resolution begin with those sequences listed at the top of their respective tables and work downwards in order to most rapidly and efficiently structurally characterize the sequence space of the conotoxins.

**Table 4:**
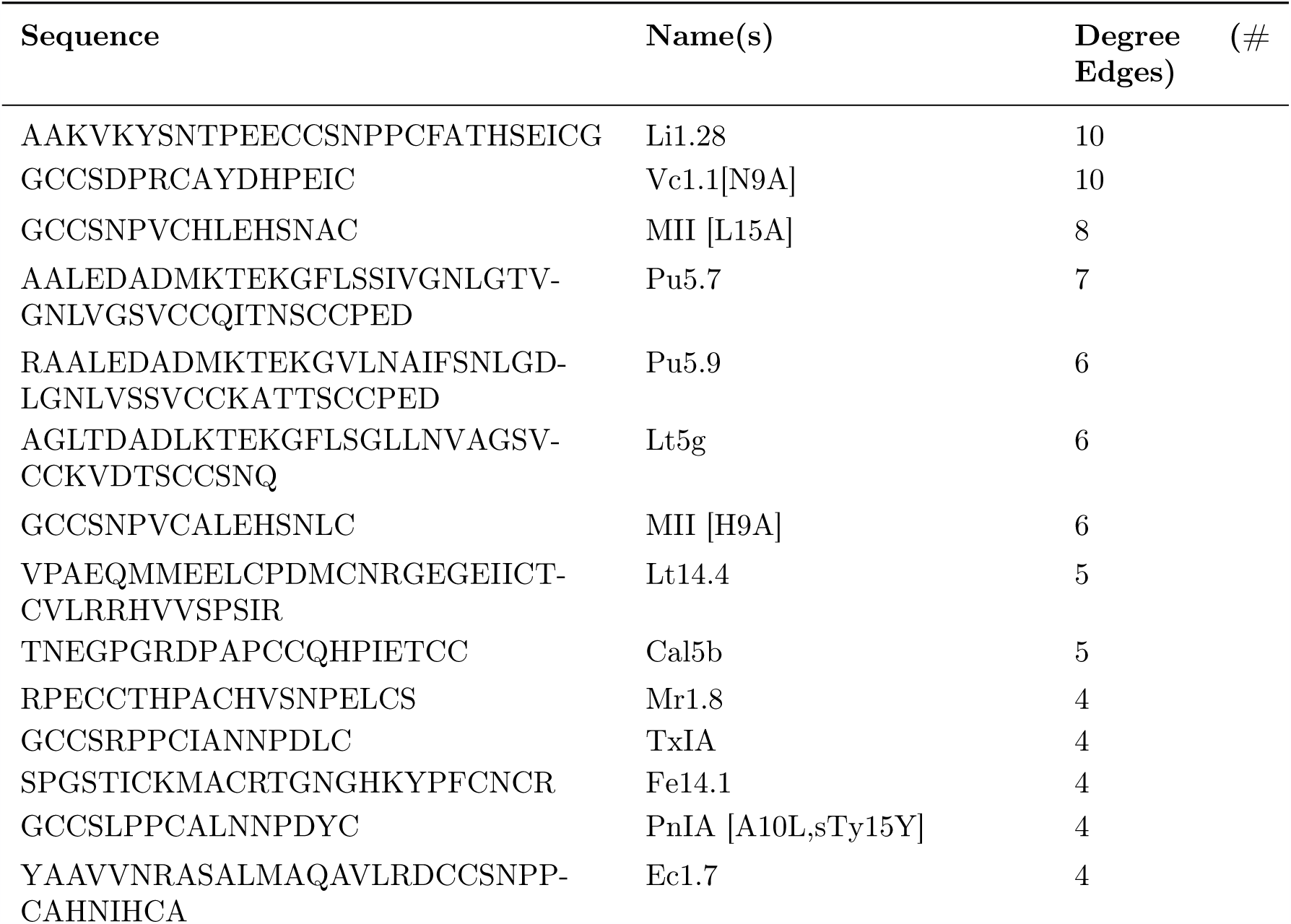
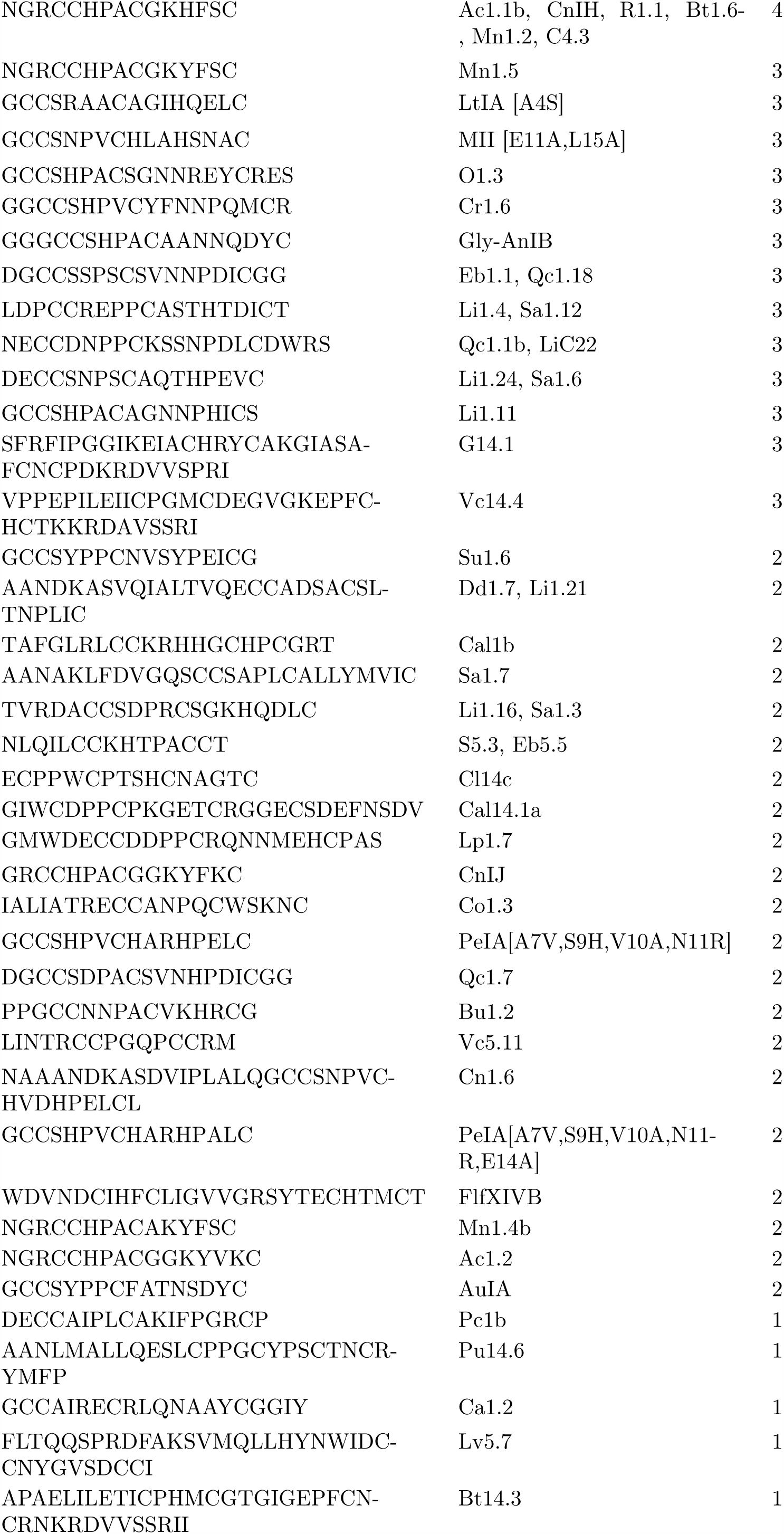
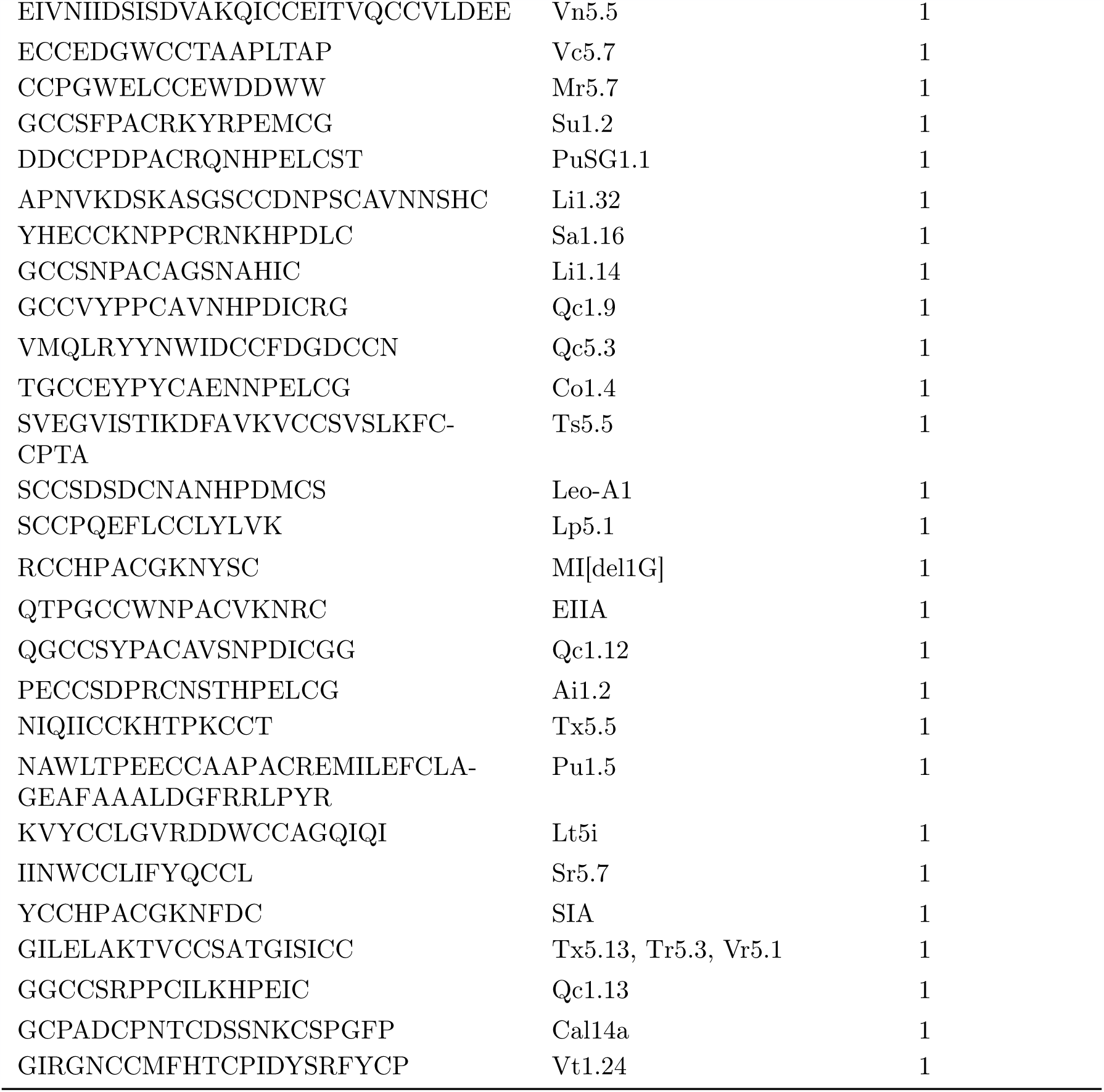
List of sequences containing four cysteines in order of interest for experimental characterization, based on degree (sequence coverage) in alignment graphs (cf. Fig. 2). Name or names of sequences are taken from the Conoserver database [Kaas et al., 2012]. Multiple names for the same sequence indicate the same sequence is produced by different species or has different post-translational modifications. Node degree corresponds to the number of sequences with pairwise alignments that are long enough and have high enough percent identity to be homology modeled with the given sequence as a template.

**Table 5:**
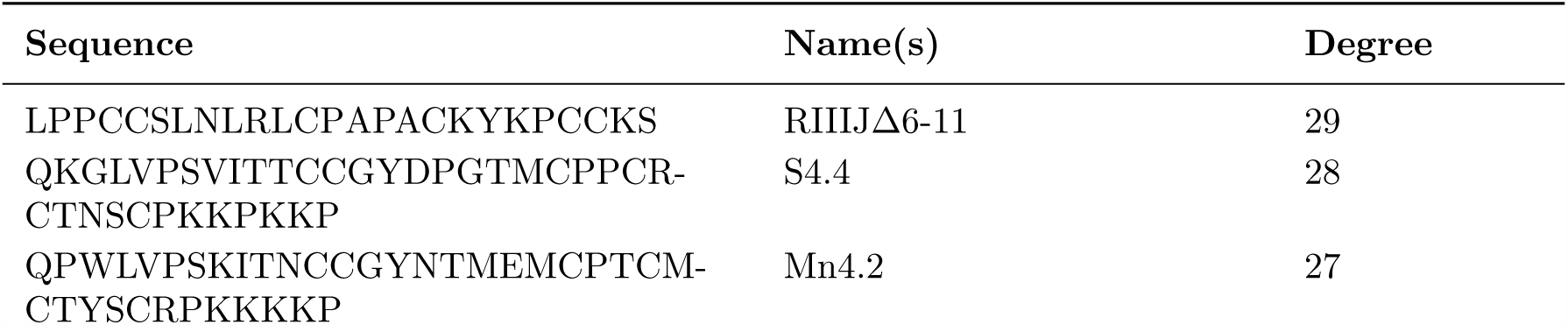
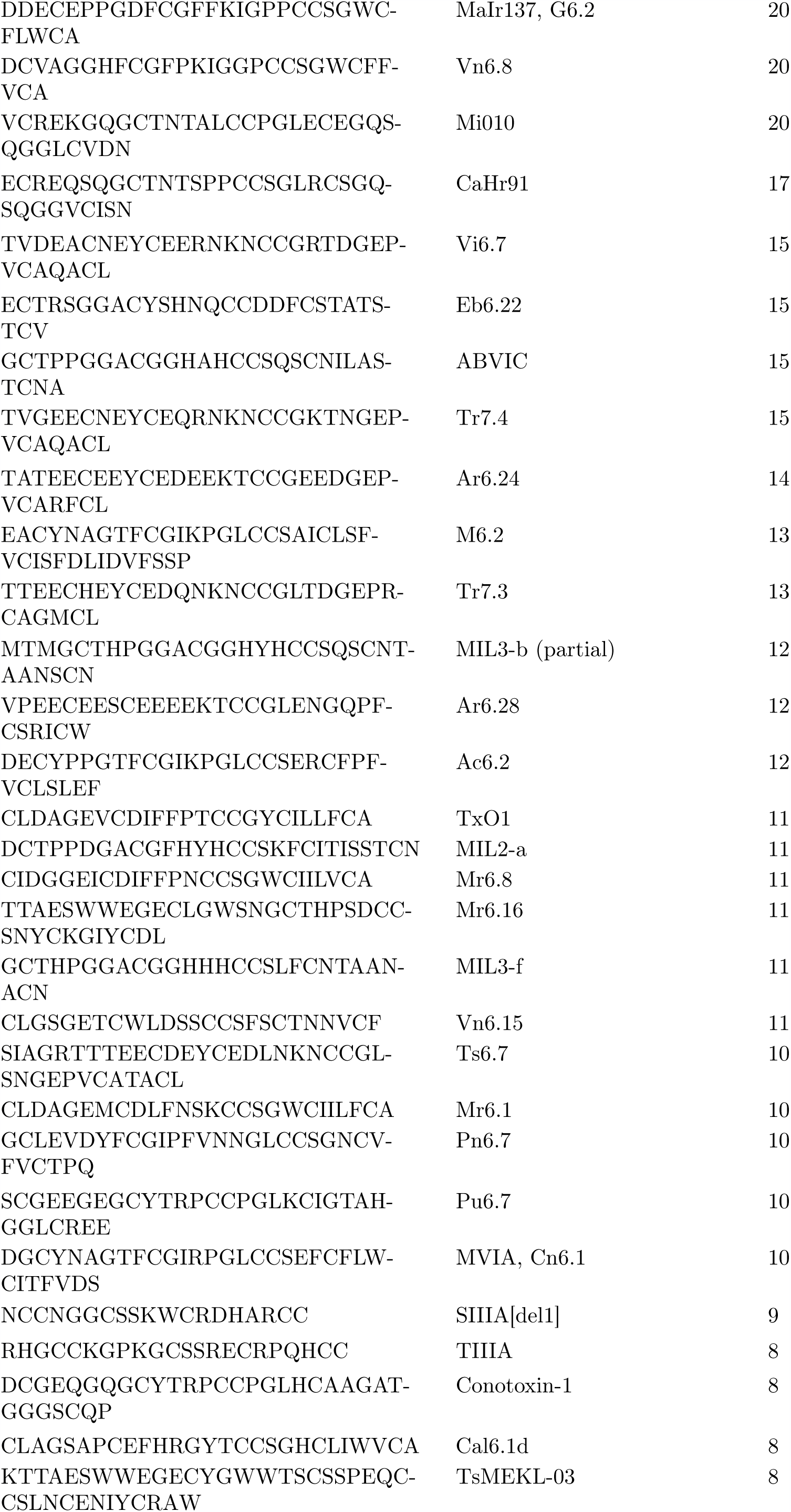
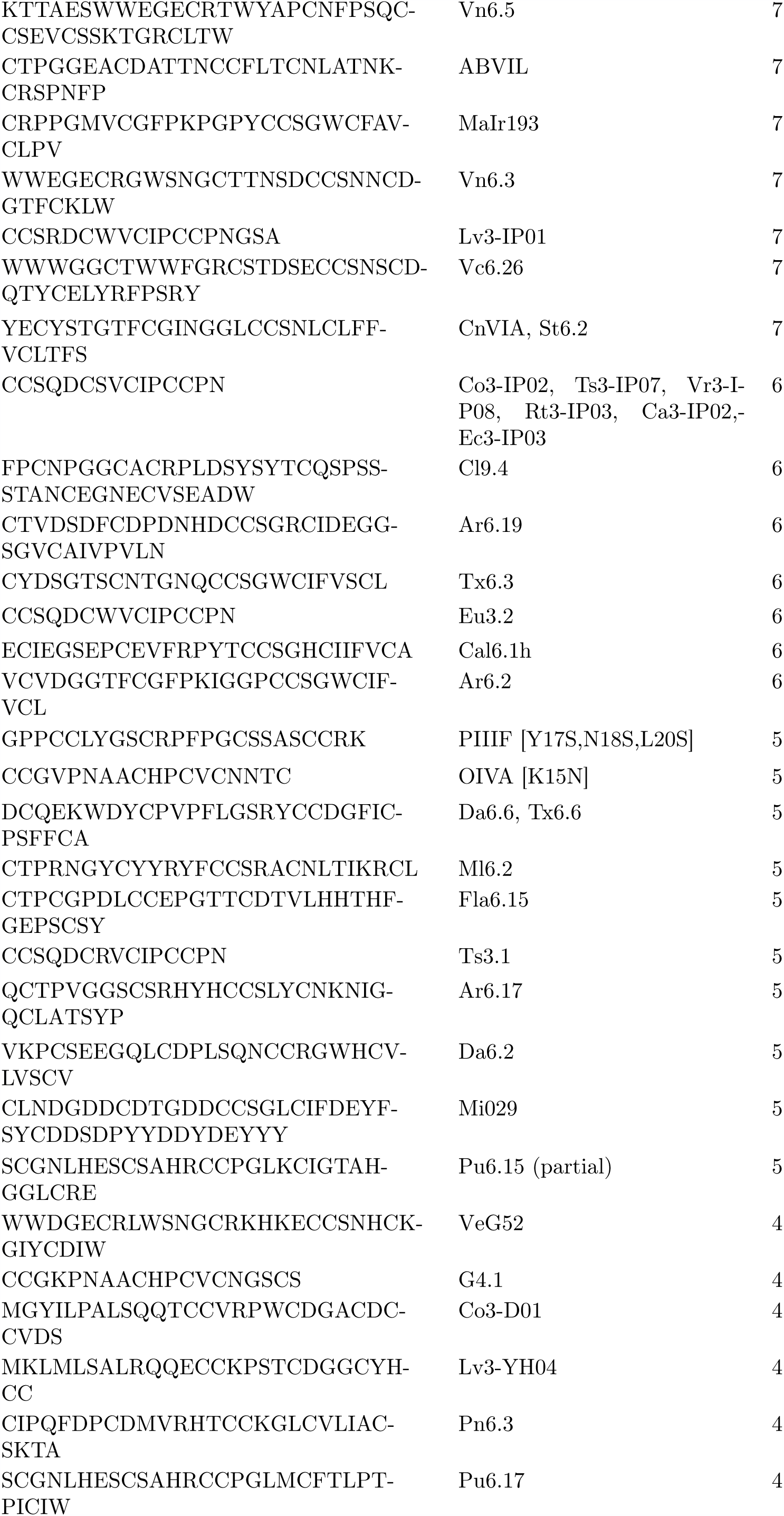
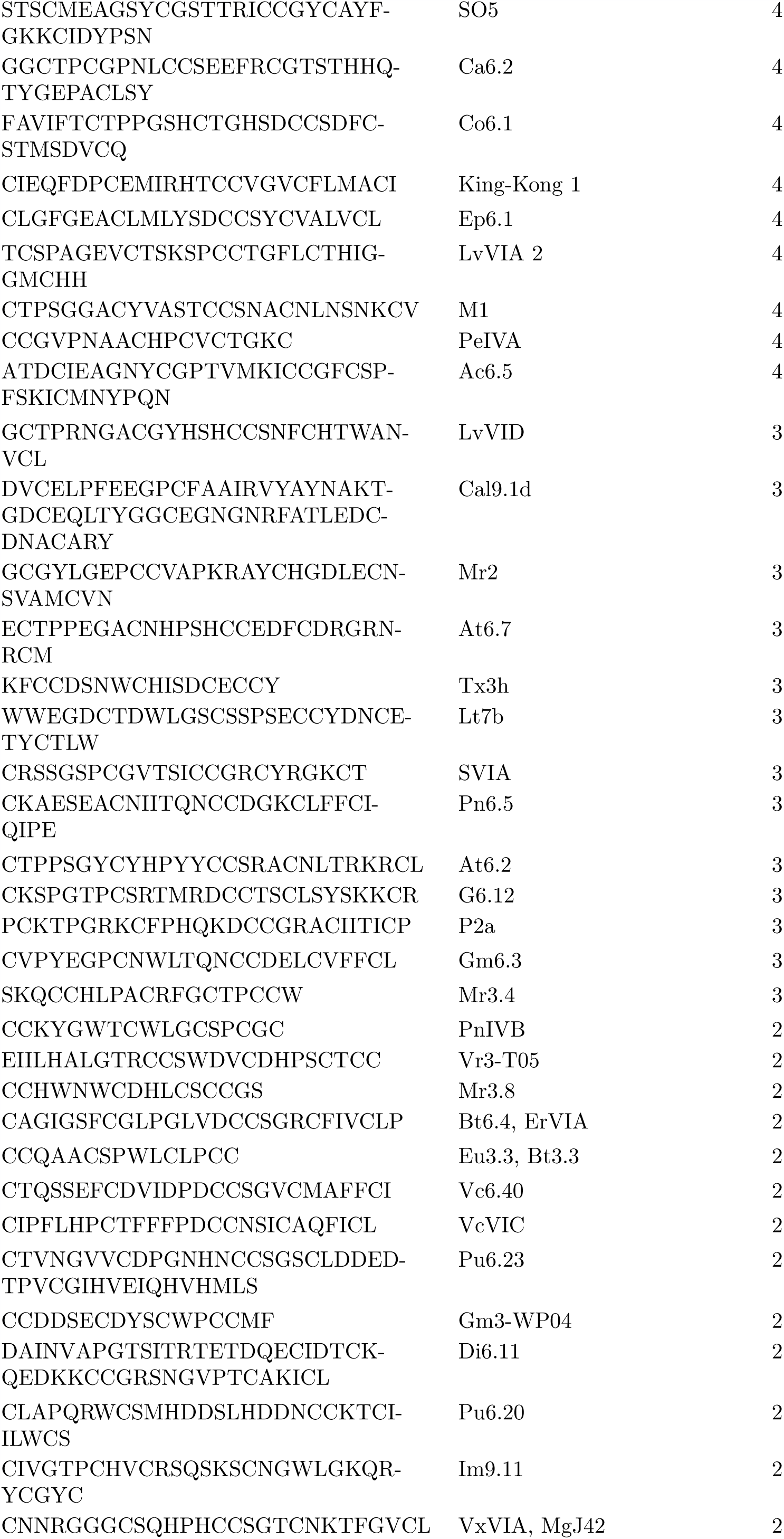
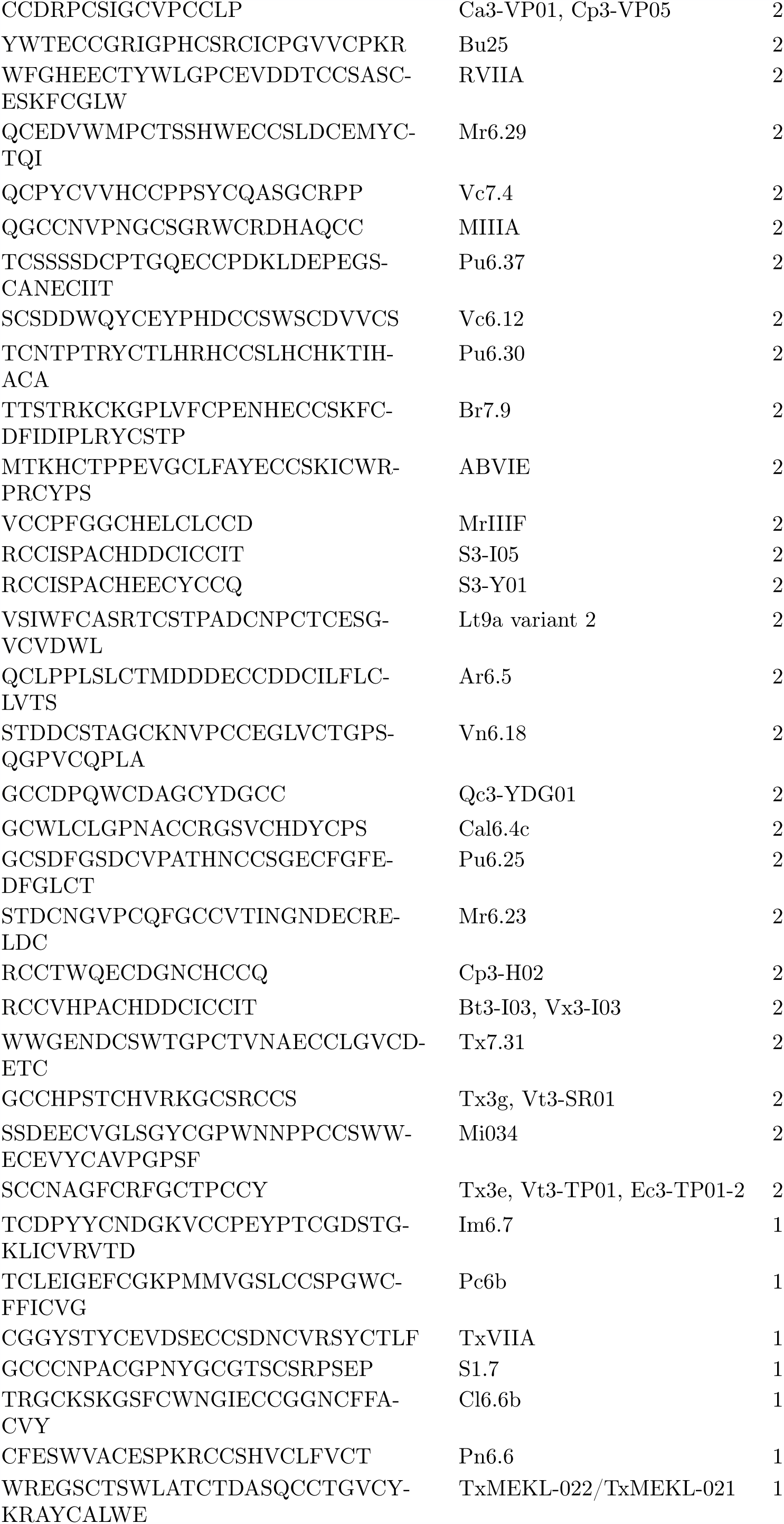
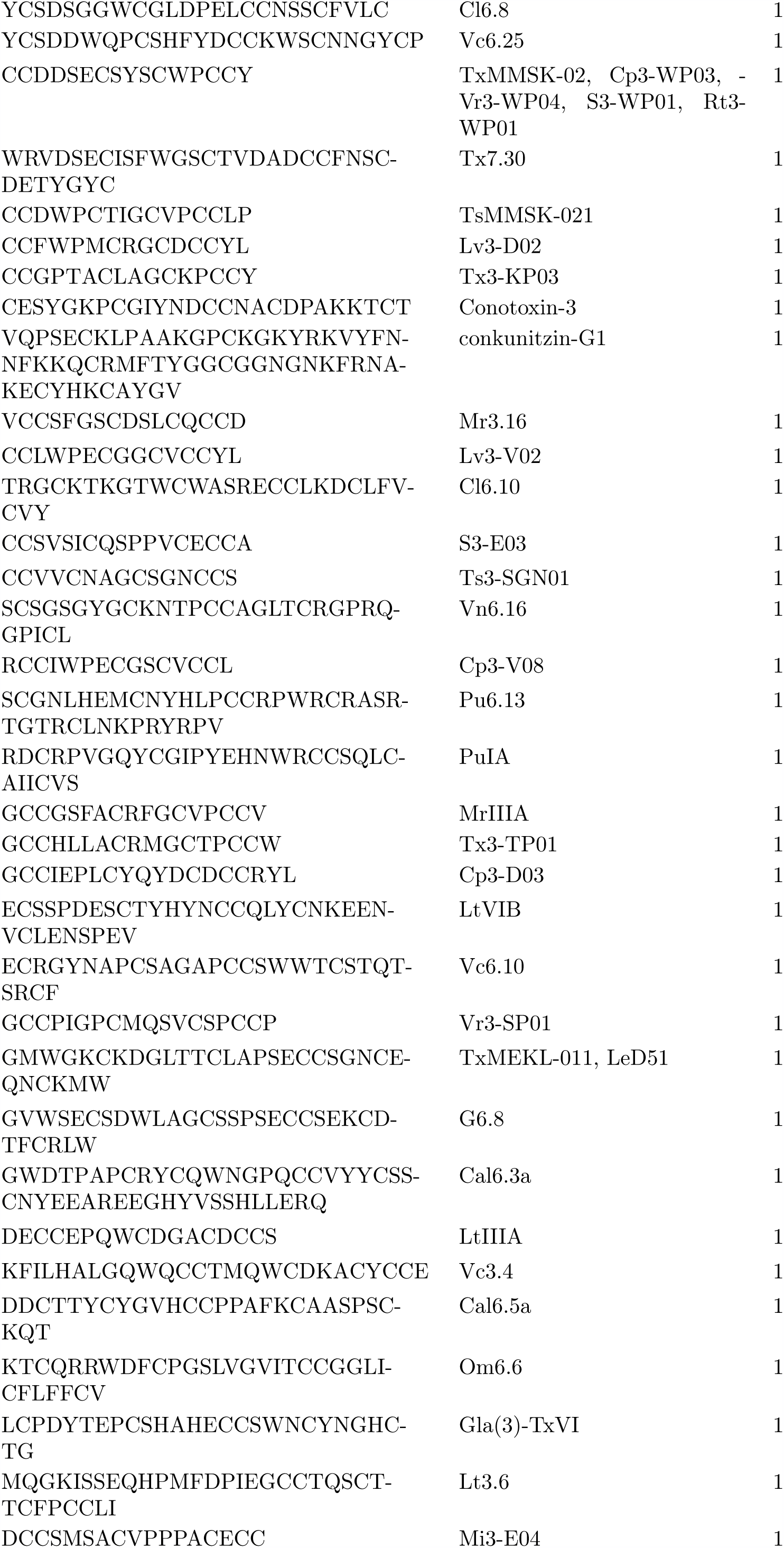
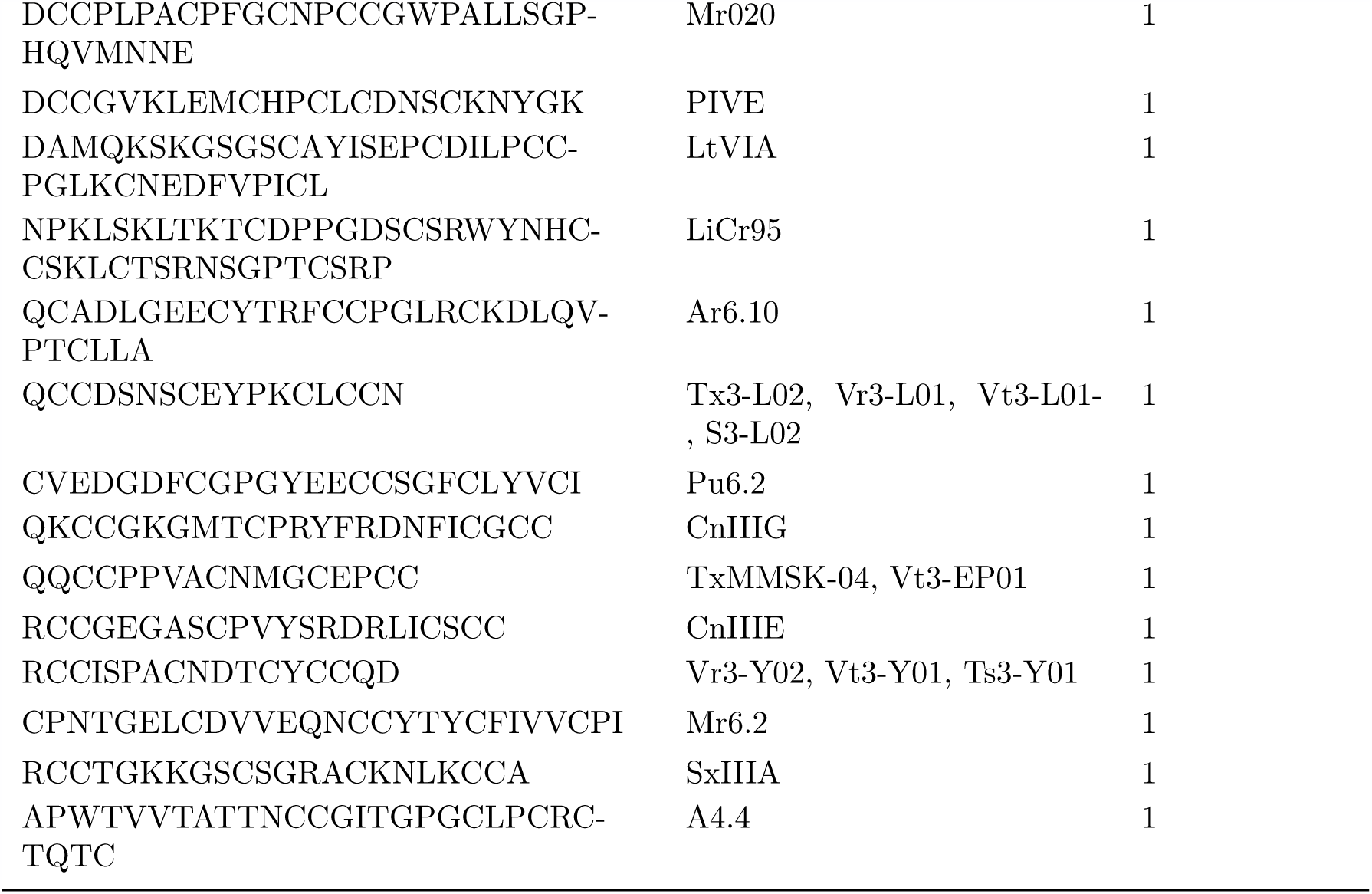
List of sequences containing six cysteines in order of interest for experimental characterization, based on degree (sequence coverage) in alignment graphs (cf. Fig. 2). Name or names of sequences are taken from the Conoserver database [Kaas et al., 2012]. Multiple names for the same sequence indicate the same sequence is produced by different species or has different post-translational modifications. Node degree corresponds to the number of sequences with pairwise alignments that are long enough and have high enough percent identity to be homology modeled with the given sequence as a template.

Finally, in Supporting File `finalmodels.zip`, we attach the set of structures computed by homology modeling, corresponding to sequences in the set {𝒞 (ℒ_ex_}), with the four, six, and eight cysteine library structures used as templates. Because we divided the sequences into subsets based on the number of cysteines in a sequence, we are able to use keeping the cysteines aligned as an additional criterion during the homology modeling procedure. The average PROCHECK G-factor, which is a log-odds score based on the likelihood of observing the given distributions of *ϕ*-*ψ* and *χ*_1_-*χ*_2_ angles in proteins, is 0.086 ± 0.005 for the reported four cysteine models, 0.10 3± 0.007 for the reported six cysteine models, and −0.2 ± 0.1 for the report eight cysteine models. Since this score is not a relative measure and values above −0.5 are generally considered acceptable, this provides evidence that the structures we have computed are physically reasonable. We further assess the quality of the homology modeling protocol by using it to model each structure in the library with templates selected from other structures in that library. The distribution of root-mean-square deviation (RMSD) values of the top three models compared with each experimental structure is shown in Fig. 4A-B. We see that our method performs well: the average RMSD in the four cysteine architecture is 2.00 ± 0.09 Å with at least 80% of the models having less than 3 Å RMSD, and the average RMSD in the six cysteine architecture is 2.3 ± 0.2 Å with 75% of the models having less than 3 Å RMSD. Most of the higher RMSD values are contributed by the flexible loops and coils. When we look at the RMSD distribution after rejecting those atoms that cannot be structurally aligned, as in case of loops and coils, the distributions improve significantly (Fig. S4), with a mean of 1.55 ± 0.09 Å for the four cysteine architecture and a mean of 1.2 ± 0.1 Å for the six cysteine architecture, with 100% of the models for both archtiectures having less than 3.5 Å deviations. A second test for validating our method was performed by checking the distribution of native contacts in the modeled structures (Fig. 4C-D). At least 60% of the native structures were captured in our models, with the distribution means of 80 ± 1% and 81 ± 1% for the four and six cysteine architectures respectively. Two pairs of residues were defined to have a native contact if the distance between the C*α* atoms in the native experimental structure was less than 8 Å, and the pair was at least 4 residues apart (C*α*^*i*^– C*α*^*i*+4^).

**Figure 4:**
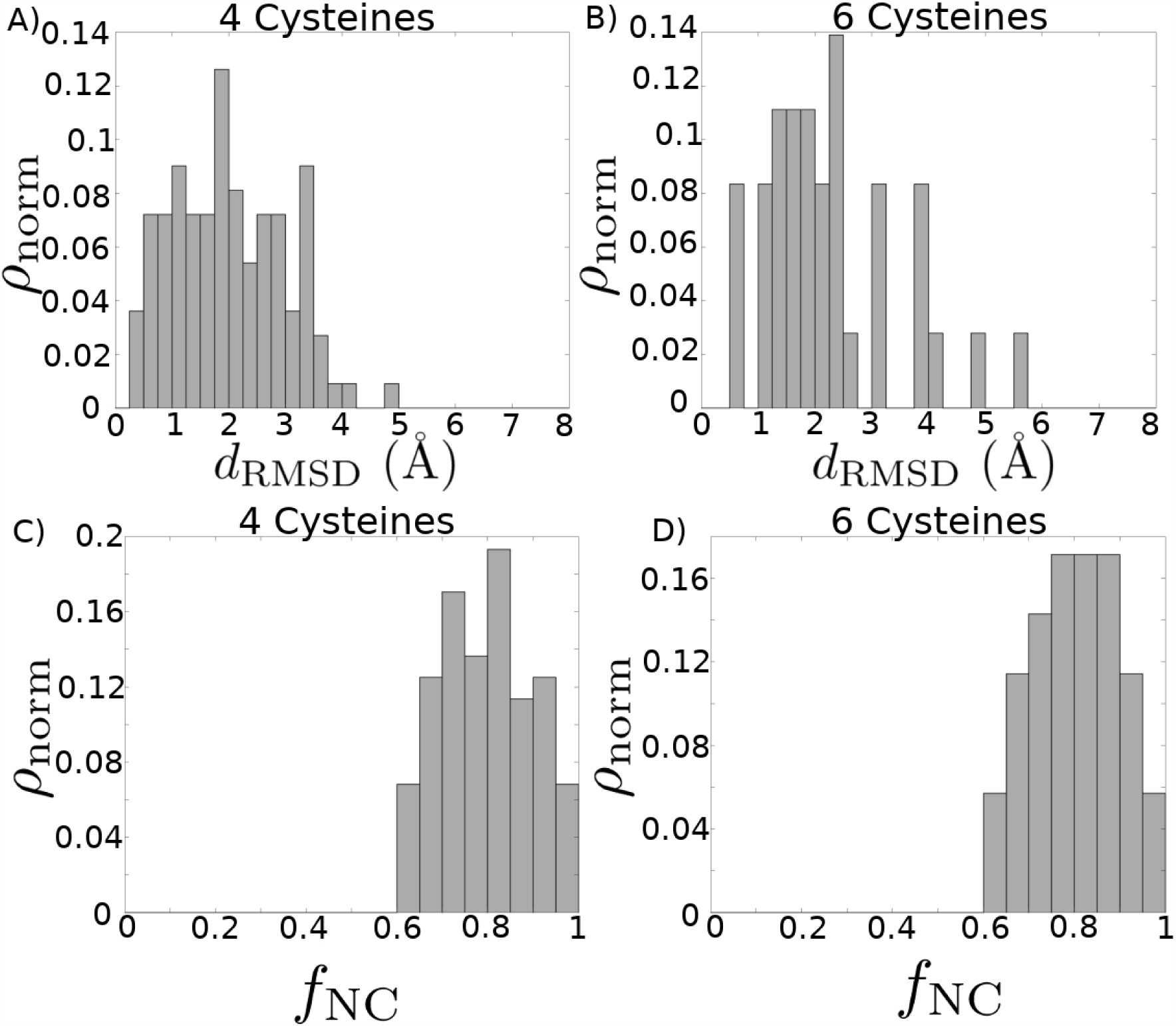
Quality of modeling criteria. (A-B) Distribution of root-mean-square deviation (RMSD) for homology models compared with their corresponding experimental structures, without prior removal of any structural alignment outliers. Each experimental structure present in the library was modeled by selecting from all other templates in the library. The top three models for each structure based on combined MODELLER DOPE and PROCHECK G-FACTOR scores are considered here. (A) Distribution mean = 2.00Å, standard deviation = 0.97Å. (B) Distribution mean = 2.25Å, standard deviation = 1.20Å. (C-D) Distribution of fraction of native contacts present in each of the homology modeled structures, with respect to the experimental structure. Each experimental structure present in the library was modeled by selecting from all other templates in the library. The top three models for each structure based on combined MODELLER DOPE and PROCHECK G-FACTOR scores are considered here. (C) Distribution mean = 0.797, standard deviation = 0.108. (D) Distribution mean = 0.805, standard deviation = 0.097.

## 3 Discussion

By employing a straightforward graph-based heuristic approach, we have constructed a set of template libraries for homology modeling of conotoxins based on the number of cysteines contained in the sequence that may also be used for homology modeling of other short, disulfide-rich, evolutionarily-related peptides. We demonstrated that libraries constructed to account for the shorter lengths of the conotoxins produce homology models that are more accurate than libraries constructed with the typical static 25% cutoff for most proteins. Currently, sufficient information is not available to homology model any sequences containing more than eight cysteines, as experimental characterization has focused preferentially on the shorter conotoxins.

Next, we employed our libraries to predict a set of structures from sequence using homology modeling, allowing us to expand the library of known conotoxin structures by about 290% overall, although a number of sequences remain without any associated structural predictions. We assessed the quality of these structures through standard techniques to demonstrate they are expected to be reasonably accurate and therefore may be employed for high-throughput screening of conotoxins as novel therapeutics for new receptor targets. In addition, our graph-based approach has allowed us to rank the remaining non-isolated sequences without corresponding characterized structures in an order that would allow for the most rapid expansion of the conotoxin structure library. We also note that of those sequences which were isolated in our graphs–that is, had no edges–80% of those containing four cysteines and 36% of those containing six cysteines were under 20 amino acids long, marking them as good candidates for a high-throughput ab initio modeling procedure, rather than necessarily for experimental characterization, as they will likely be tractable but will not contain any information about other sequences.

One important point about short, disulfide-rich peptides that we have not addressed in this work is the existence of so-called “disulfide isomers.” Under certain conditions, there is experimental evidence suggesting that some toxins do not exist as a single set of “native” structures but as a heterogeneous–perhaps metastable–ensemble populated with strikingly different secondary structures corresponding to differing patterns of cysteine connectivity [Paul George et al., 2018, Combelles et al., 2008]. Characterizing multiple possible disulfide isomers is outside the purview of homology modeling, but it is an important area of future work and sounds a note of caution on the standard interpretation of structure libraries, which generally assume a single “native” structure dictated wholly or primarily by the folding propensities of the amino acid sequence.

Overall, the work in this article presents a rational graph-based algorithm that we employ to expand the repertoire of known conotoxin structures for application in a high-throughput manner as part of the early stages of drug design. We expect that the libraries, the expanded set of structures, and the ranking of sequences in terms of degree of connectedness to other sequences will be valuable resources improving the prospects of conotoxins as novel therapeutic leads and that our approach may be employed for further characterization of other sets of evolutionarily-related toxins.

## Supporting information

Template Libraries

Homology Modeled Structures

## 4 Acknowledgments

T.T and S.G. were supported by the Functional Genomic and Computational Assessment of Threats (Fun-GCAT) program of the Intelligence Advanced Research Projects Activity (IARPA) agency within the Office of the Director of National Intelligence. S.C. and T.T. were also partially supported by the Center for Nonlinear Sciences (CNLS) at LANL. R.A.M gratefully acknowledges a Los Alamos National Laboratory Director’s Postdoctoral Fellowship. Triad National Security, LLC (Los Alamos, NM, USA) operator of the Los Alamos National Laboratory under Contract No. 89233218CNA000001 with the U.S. Department of Energy. We thank Dr. Will Fischer for valuable discussions. This research used resources provided by the Los Alamos National Laboratory Institutional Computing Program, which is supported by the U.S. Department of Energy National Nuclear Security Administration under Contract No. 89233218CNA000001.The views and conclusions contained herein are those of the authors, and should not be interpreted as necessarily representing the official policies or endorsements, either expressed or implied, of the Office of the Director of National Intelligence (ODNI), Intelligence Advanced Research Projects Agency (IARPA), Los Alamos National Laboratory (LANL), Department of Energy (DOE), or the US Government.

## 5 Author Contributions

Conceptualization, R.A.M., S.C., T.T., and S.G.; Methodology, R.A.M., S.C., T.T., and S.G.; Software, R.A.M., S.C., and T.T.; Formal Analysis, R.A.M., S.C. and T.T.; Investigation, R.A.M., S.C. and T.T.; Writing – Original Draft, R.A.M.; Writing – Review & Editing, R.A.M., S.C., T.T., and S.G.; Visualization, R.A.M. and S.C.; Supervision, S.G.; Project Administration, S.G.; Funding Acquisition, S.G.

## 6 Declaration of Interest

The authors declare no competing interests.

## 7 STAR Methods

### 7.1 Contact for Reagent and Resource Sharing

Further information and requests for resources and reagents should be directed to and will be fulfilled by the Lead Contact, S. Gnanakaran

### 7.2 Method Details

For use in construction of the template libraries, we employed a set of 142 conotoxin structures downloaded from the PDB [Berman et al., 2000], which we found by searching “conotoxin” on the PDB. We manually removed several false positives, such as a crystal structure of the acetylcholine-binding protein that was identified due to the title of the associated paper. We also manually removed several sequences that were identical to natural conotoxin sequences but modified by the replacement of disulfide bonds with dicarba bonds. We did not remove redundant sequences consisting of multiple characterization methods and in a few cases structural isomers resulting from different disulfide-bond connections. In future, further work will be done to properly assess the likelihood of multiple stable or metastable states, but we do not address this consideration further here.

For use in the analysis detailed in this article, we downloaded a set of 6,255 peptide sequences from the Conoserver [Kaas et al., 2012] using the Tools > Download Conoserver’s Data command. We retained only sequences containing four, six, eight, or ten cysteines. We removed anything with the word “precursor” or “patent” in the name, as precursor sequences contain, in addition to the mature peptide sequence that folds into the toxin, a signal sequence and N-and C-terminal pro-regions that are cleaved in the endoplasmic reticulum and Golgi apparatus [Kaas et al., 2010]. A manual inspection of sequences labeled “patent” revealed that many were insufficiently characterized–for example, they noted only the cysteine pattern or they mixed precursor and mature toxin sequences with no indication. We also added to the sequence list any sequence that corresponded to one of the PDB structures that was not already contained in the list. Once the set of all sequences was finalized, we split it into four subsets corresponding to the number of cysteines contained. In the end we retained for analysis a total of 801 unique sequences containing four cysteines, 1,113 unique sequences containing six cysteines, 190 unique sequences containing eight cysteines, and 53 unique sequences containing ten cysteines.

#### 7.2.1 Library template selection procedure

For each subset of sequences corresponding to a different number of contained cysteines, we created an alignment graph as follows. For every sequence we computed a pairwise alignment with every other sequence, using the `PairwiseAligner` class in the `Align` module of the Biopython package [Cock et al., 2009], in global mode, with a gap-open penalty of −10 and a gap-extend penalty of −0.5. Employing the `networkx` Python package [Hagberg et al., 2008], we constructed a graph in which nodes represented sequences and we placed an edge between two nodes whenever the percent identity of the highest-percentage pairwise alignment of the two corresponding sequences was greater than [Rost, 1999],

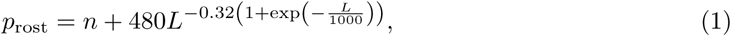

where *L* is the length of the alignment in numbers of amino acid residues and we set *n* = 5 (%).

We constructed two different template libraries for each subset of sequences, one from the pairwise alignment graph and one from a static 25%-identity cutoff (with *n* = 5 %). When creating the graph-based libraries (see also Fig. 1), we first identified all connected components in the graph. For each connected component, we chose first the sequence with the highest node degree (number of distinct edges) that corresponded to a structure in the set of 142 structures downloaded from the PDB, added it to the library, and removed that sequence and all sequences it shared an edge with from the graph. We continued this procedure until one of two criteria was satisfied: (i) there were no longer any sequences in the connected component with corresponding structures or (ii) there were no longer any sequences in the connected component without corresponding structures. These criteria corresponded to the following two situations, respectively: (i) there were no other structures available for inclusion in the library or (ii) the entire connected component was able to be homology modeled based on the structures included in the library up until that point. For construction of the static sequence-identity cutoff library, sequences within each data set were clustered and a representative sequence from each cluster chosen by using the `sequence_db.filter` command of MODELLER version 9.20 [Sali and Blundell, 1993, Webb and Sali, 2016], which groups sequences together if their sequence identity is greater than a specified cutoff value. The set of cluster representatives became a library of structures in which between any pair the sequence identity was less than the specified cutoff value.

For computation of the homology modeled structures based on the library templates that were used to assess and compare the quality of the two libraries, we used the align2d command followed by the automodel procedure from Modeller 9.20 with default parameters. We computed five models for each sequence from each template (except for itself, in the case of library structures being modelled based on other library structures). The best homology model was chosen as the one with the lowest backbone RMSD to the known or experimentally-resolved structure, using the align command in PyMOL [Schrödinger LLC, 2015] that superimposes two structures via a structure superposition that is constrained by a prior sequence alignment.

#### 7.2.2 Homology modeling criteria

After assessing the quality of the template libraries, we used the graph-based libraries to construct via homology modeling a database of structures for those conotoxin sequences (set {𝒞 (ℒ_ex_}) shown in blue in Figs. 2 and S2) that are covered by those libraries (set {ℒ_ex_} given in orange in Figs. 2 and S2). The pipeline employed for building these homology-modeled structures is detailed here. A schematic of this pipeline is given in Fig. 5. The 4C subset included 143 such sequences for which structures were computed by homology modeling from 49 library structure templates, while the 6C subset included 148 sequences for which structures were computed by homology modeling from 30 library structure templates. There was only one existing non-isolated library structure template having 8C architecture, and 17 sequences were modeled from it, while no 10C structures could be modeled, since to date there are no structures of conotoxins containing 10 cysteines deposited in the PDB.

**Figure 5:**
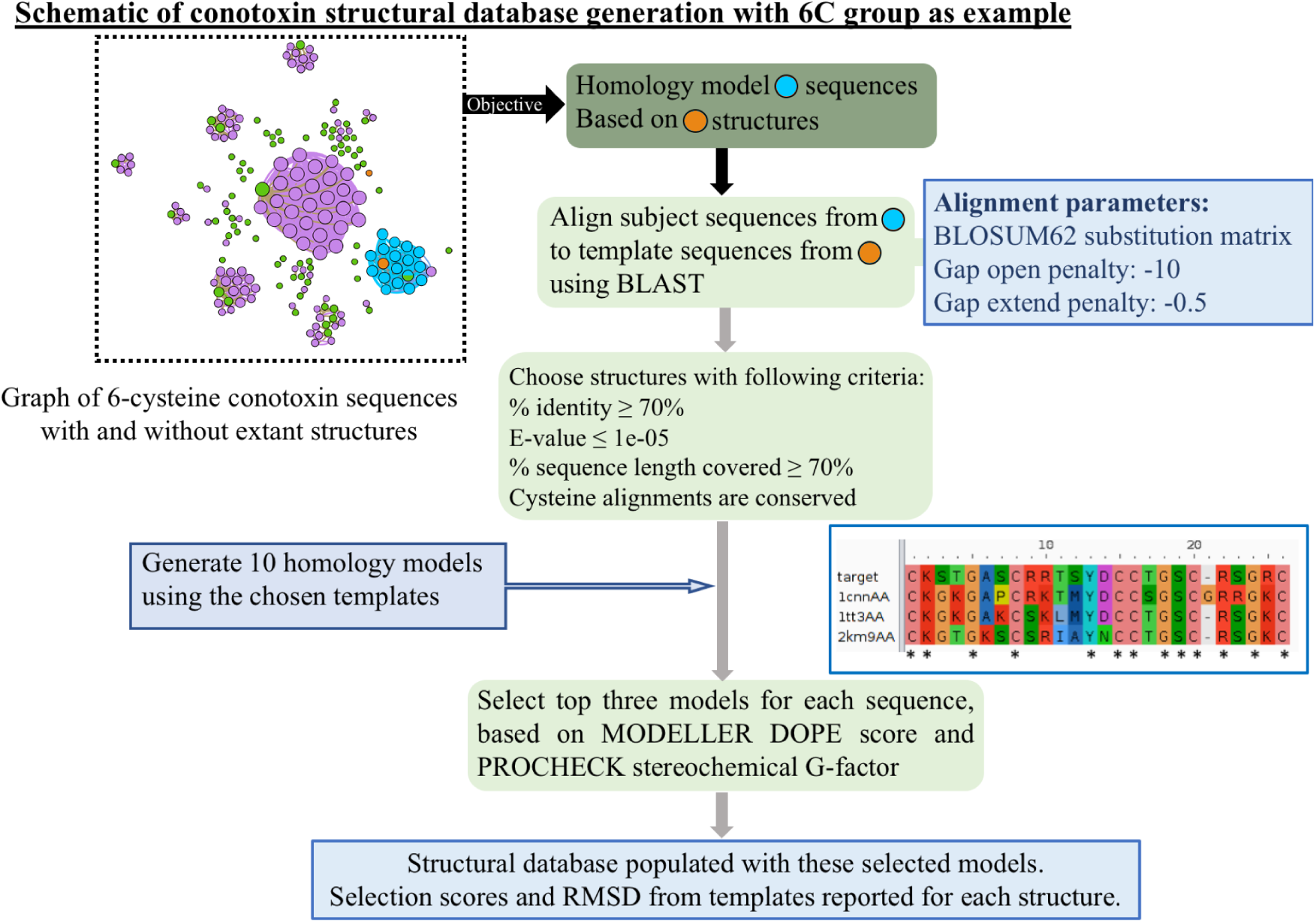
Schematic of procedure for producing homology modeled structures from library templates for conotoxin sequences with unknown structure lying in the set {𝒞 (ℒ_ex_}. Graph inset of eight cysteine graph is an example. The inset consisting of an example alignment input figure was created using the alignment obtained from BLAST [Altschul et al., 1990] and visualized with Aliview [Larsson, 2014].

Alignment of each of the subject sequences was performed with those sequences that have a structure present in the template library using BLAST [Altschul et al., 1990]. BLOSUM62 substitution matrix [Henikoff and Henikoff, 1992] was used with a gap-open penalty of −10 and a gap-extend penalty of −1. For each sequence, structures were considered possible templates if they fulfilled the following criteria: (i) sequence identity of ≥ 70%; (ii) ≥ 70% of sequence length covered; (iii) E-value ≤ 1 × 10^−5^. Additionally, we constrained the cysteines in the sequence to be aligned in the following manner. If there was a one-position shift in the sequence alignment that would allow the cysteines to align, the gap penalties at that position were removed to enforce cysteine alignment. If a greater than one-position shift would be required to allow the cysteines to align, such a template was not considered.

Structural homology modeling was performed using the MODELLER version 9.20 package [Sali and Blundell, 1993, Webb and Sali, 2016]. Multiple templates were used to aid in the modelling process for those subjects where more than one sequence satisfied the above-mentioned criteria. The models were further relaxed by several steps of conjugate gradients and molecular dynamics with simulated annealing as recommended in the thorough Variable Target Function Method (VTFM) optimization of MODELLER [Braun and Go, 1985]. Due to the alignment of cysteines from the template structures, the disulfide bonds could be constrained by patches. Ten such models were generated for each subject sequence. Subjects 107 and 110 from 4C architecture and subject 2 from 6C architecture did not correspond to any templates that satisfied all of our above criteria. Nevertheless, we modeled these sequences based on the best sequence match.

We selected three top models for each subject based on the Discrete Optimized Protein Energy (DOPE) score [Shen and Sali, 2006] and the PROCHECK G-factor [Laskowski et al., 1993]. DOPE, a typical criterion for assessing the quality of a modeled structure, is an atomistic distance-dependent statistical potential calculated from a large set of refined high resolution PDB structures. The PROCHECK G-factor is a log-odds score based on observed distributions of the *ϕ*-*ψ*, and *χ*_1_-*χ*_2_ values measuring whether the model is physically reasonable or if it contains unusual stereochemical configurations. In this study, we normalized the DOPE and G-factor scores and used a combined product of probabilities to sort the structures. The top three models selected for each subject are reported in Supplementary File `finalmodels.zip`, along with their DOPE, G-factor, MODELLER optimization function value (molpdf), GA341 scores [John and Sali, 2003], and the Ramachandran plots for each of these models. All RMSD calculations were performed with Pymol [Schrödinger LLC, 2015]. There is only one available non-isolated structure in the 8C extant library. This was used to model all 17 subject sequences. The best three models for each sequence along with their assessment scores are reported in the database.

### 7.3 Quantification and Statistical Analysis

We use the Kolmogorov-Smirnov two-tailed test as implemented in the SciPy package [Oliphant, 2007] and referred to by the `ks2samp` command to assess whether we may reject the null hypothesis of the RMSDs of experimental structures from homology models based on different template libraries being drawn from the same distribution. We employ a significance level of *p* = 0.05, meaning that we reject the null hypothesis if the KS statistic *D* returned by the test is such that 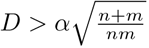, where *α* = 1.224 for a significance level of *p* = 0.05, and *n* and *m* are the number of samples in each set respectively. The analysis is referred to in Sec. 2.

### 7.4 Data and Software Availability

Data is available as supplementary files and software is available upon request from the corresponding author (see Next Section).

### 7.5 Supplemental Information

We provide tables of 8C and 10C sequences in order of projected interest for experimental characterization and four supplementary figures. Structures used as template libraries are provided in the supplementary data folder `libraries.zip`. Homology modeled structures of conotoxins are provided in the supplementary data folder `finalmodels.zip`, along with their scores and the associated Ramachandran plots. Python, Modeller, and Bash analysis scripts for preparation, graph construction and further analysis will be provided upon request.

## A Supplemental Information

**Figure S1:**
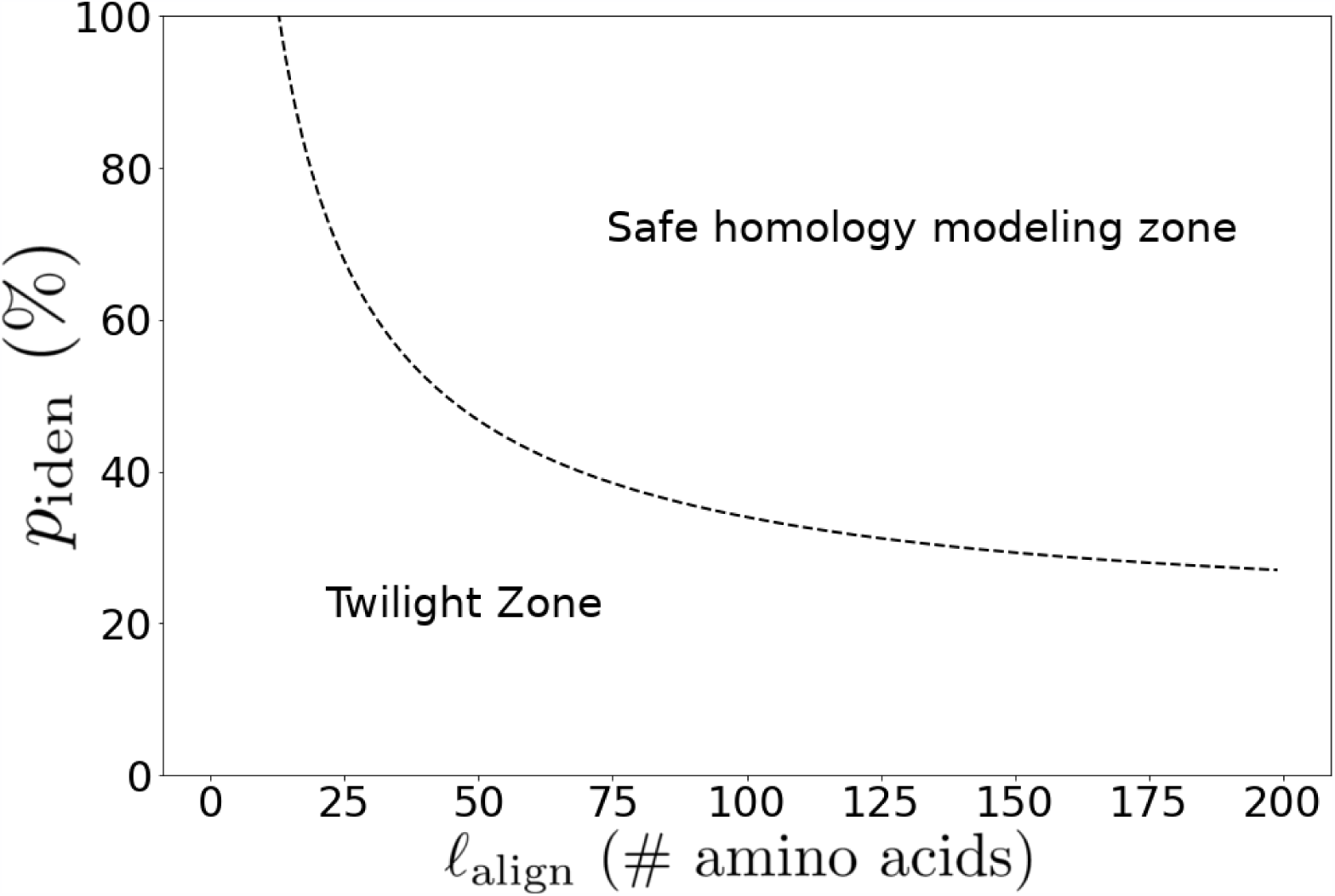
Rost’s phenomenological curve (Eqn. 1) of minimum percentage identity for homology modeling as a function of pairwise alignment length with *n* = 5% padding as employed in this work. As the length of the alignment decreases, the minimum percent identity for homology modeling increases, and there is a particularly rapid increase below alignments of about 25 amino acids, where a fairly large proportion of toxins reside.

**Table S1:**
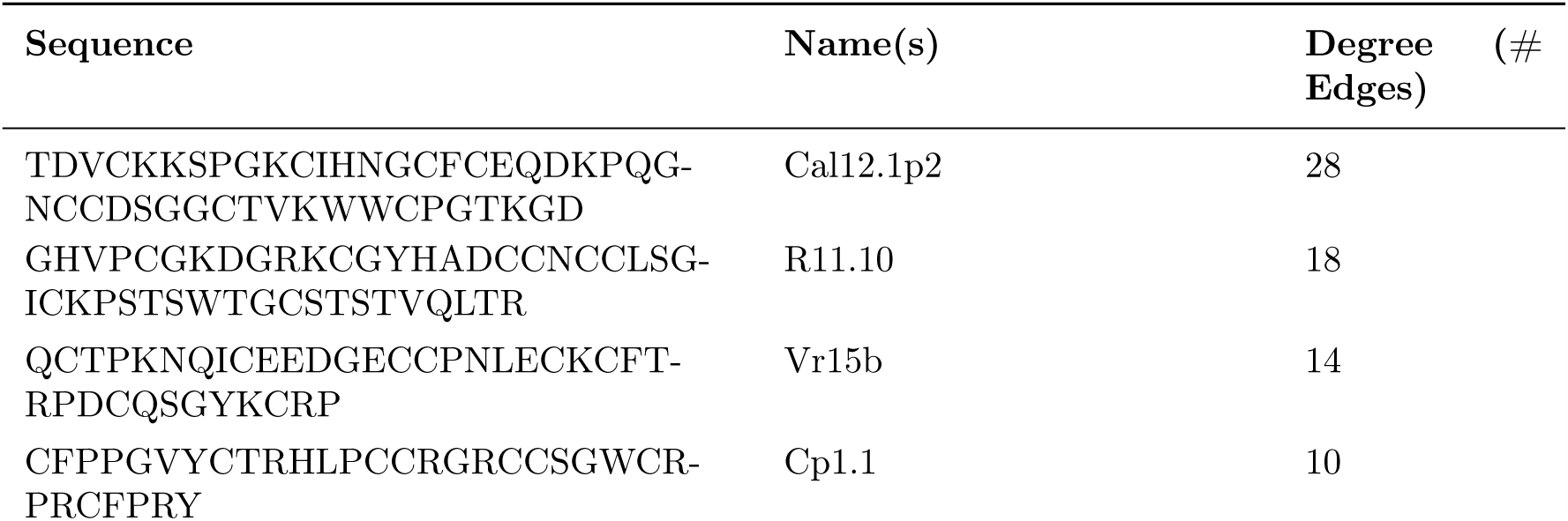
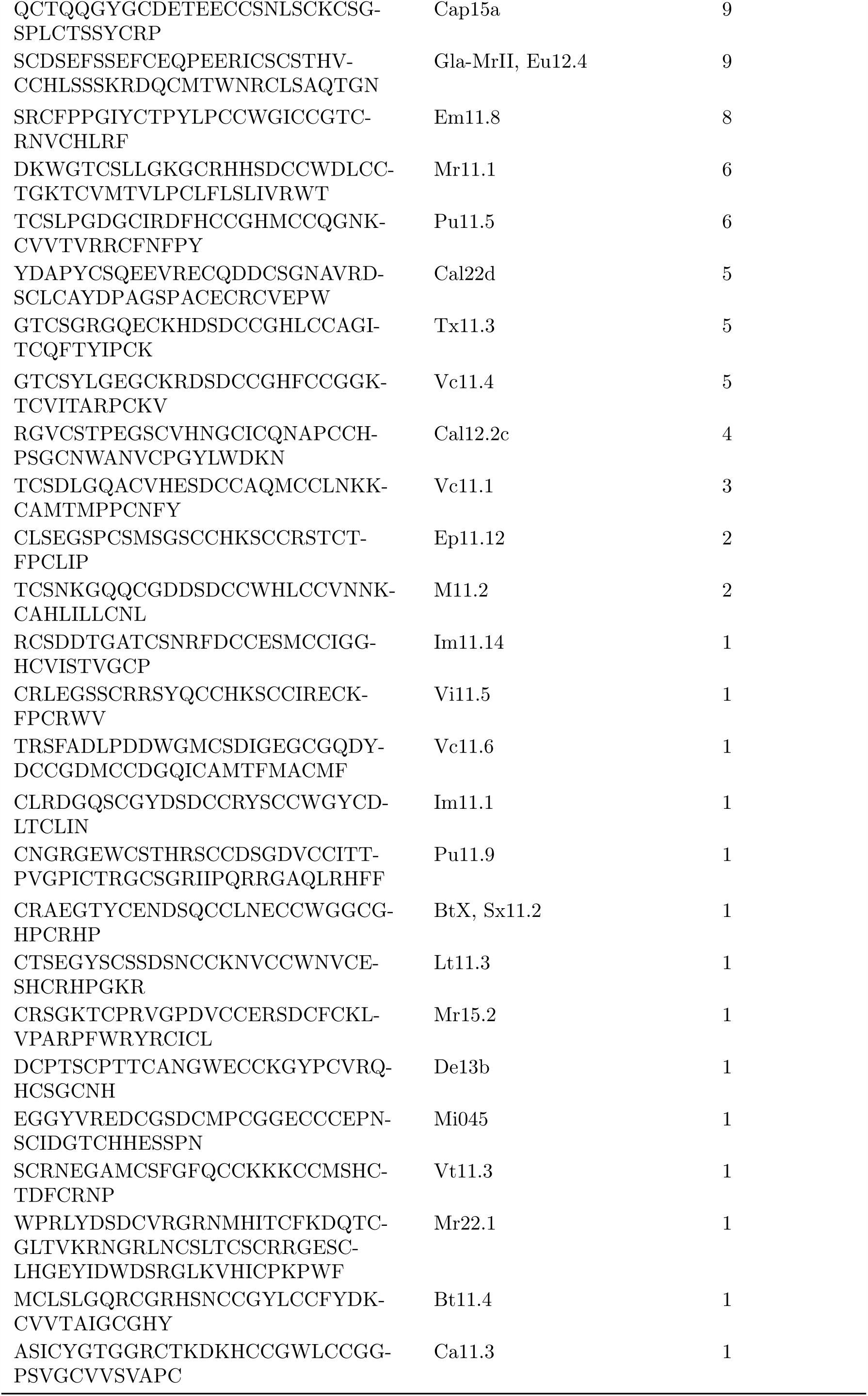
List of sequences containing eight cysteines in order of interest for experimental characterization, based on degree (sequence coverage) in alignment graphs (cf. Fig. 2). Name or names of sequences are taken from the Conoserver database [Kaas et al., 2012]. Multiple names for the same sequence indicate the same sequence is produced by different species. Node degree corresponds to the number of sequences with pairwise alignments that are long enough and have high enough percent identity to be homology modeled with the given sequence as a template.

**Figure S2:**
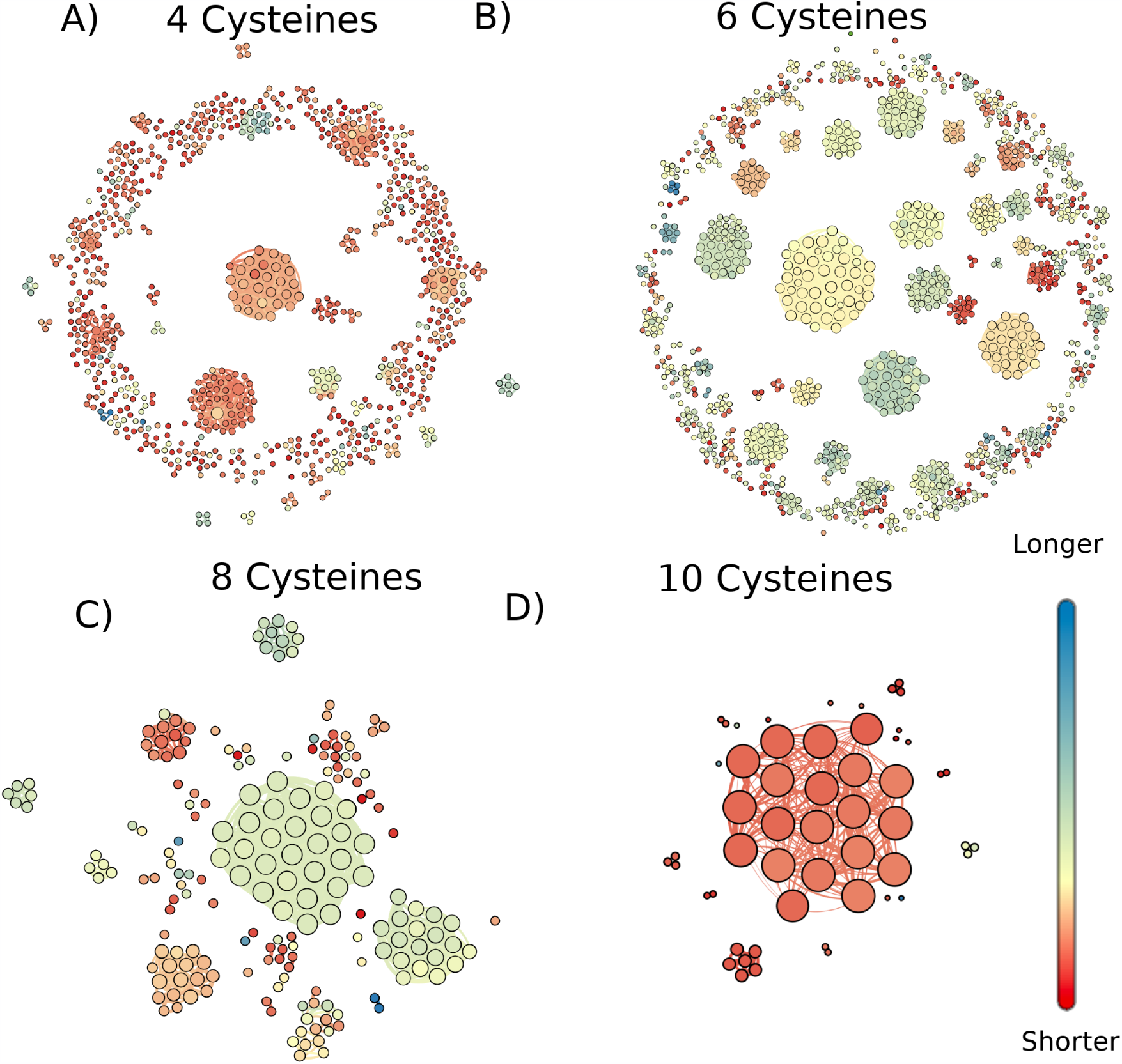
Graph of conotoxins containing (A) four cysteines, (B) six cysteines, (C) eight cysteines and (D) ten cysteines where nodes are sequences and edges exist between sequences with pairwise alignments that have high enough length and percent identity to fall above the Rost curve with *n* = 5% (Eqn. 1). Colors show the relative sequence lengths of each graph, but the color scale of each graph is independent of the others. The sizes of the nodes corresponds to their degree; that is the number of other sequences that they can be modeled based on or used to model. Node locations and edge lengths were chosen for ease of visualization of separate connectec components. Visualization of the graphs was produced with Gephi 0.9.2 [Bastian et al., 2009].

**Table S2:**
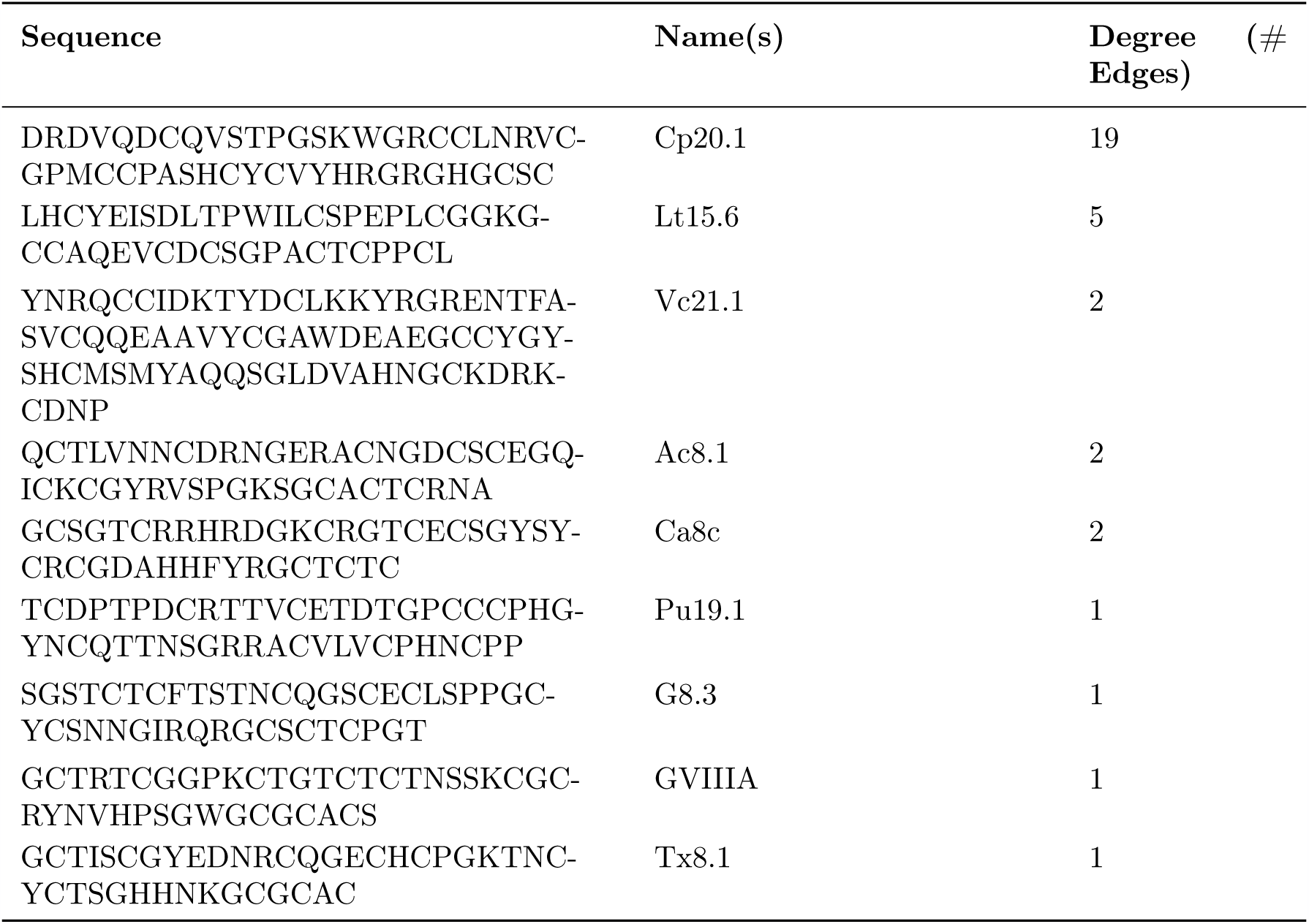
List of sequences containing ten cysteines in order of interest for experimental characterization, based on degree (sequence coverage) in alignment graphs (cf. Fig. 2) Name of sequences are taken from the Conoserver database [Kaas et al., 2012]. Node degree corresponds to the number of sequences with pairwise alignments that are long enough and have high enough percent identity to be homology modeled with the given sequence as a template.

**Figure S3:**
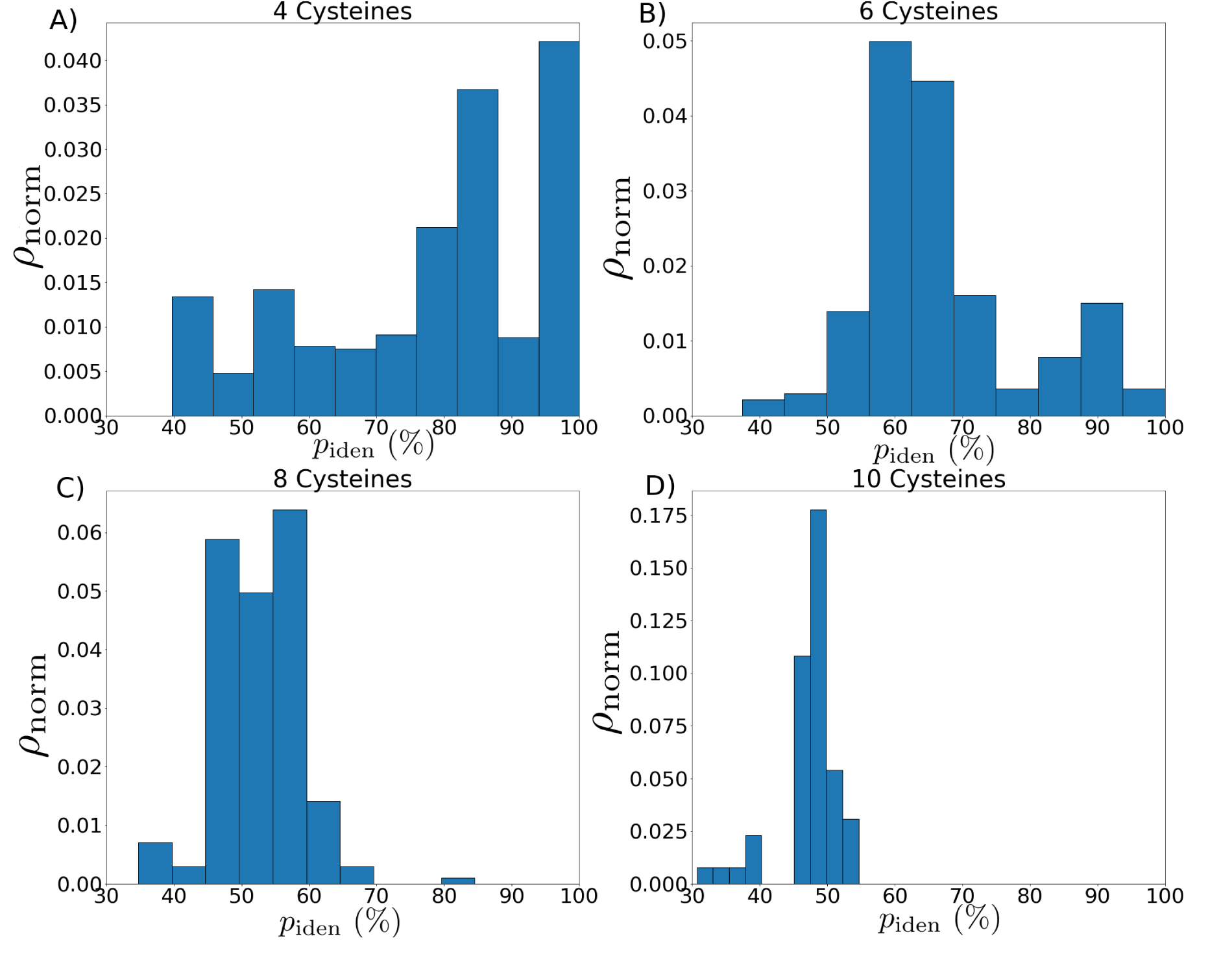
Distributions of approximate minimum percentage identity cutoffs for the conotoxins containing A) four, B) six, C) eight, and D) ten cysteines. We assume for demonstration purposes that any alignment is the length of the peptide itself. We employ Rost’s curve (see Eqn. 1 and Fig. S1) with a padding of *n* = 5%. The distribution shifts significantly downward as the number of cysteines and concomitantly the overall length of the peptides under consideration increases. Note too the presence in panels A) and B) of a bin going up to 100%, which demonstrates the existence of peptides so short among the conotoxins that it is impossible to reliably predict their structure via homology modeling, which comprise a large proportion of the isolated nodes in the graphs (cf. Fig. S2)

**Figure S4:**
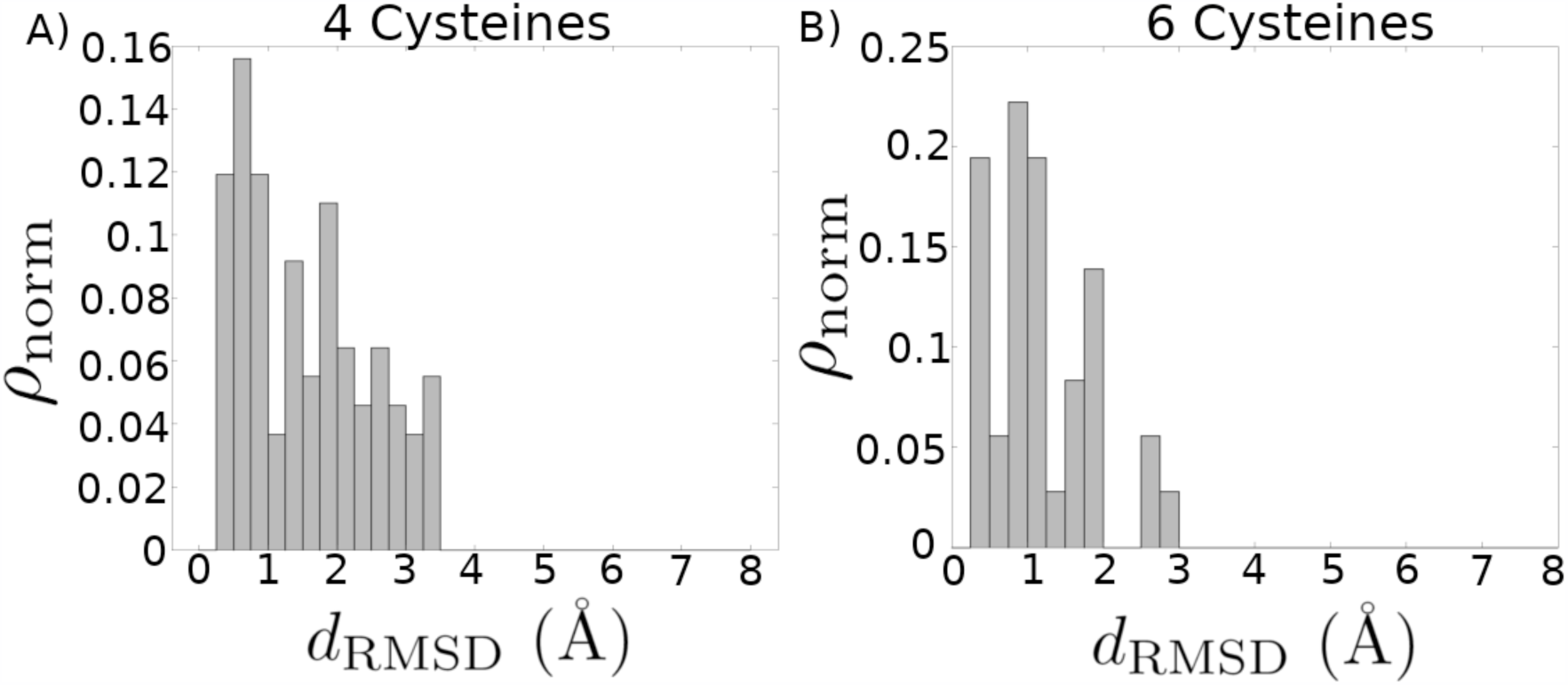
Distribution of root-mean-square deviation (RMSD) for homology models compared with their corresponding experimental structures, after refinement involving rejection of structural alignment outliers. Each experimental structure present in the library was modeled by selecting from all other templates in the library. The top three models for each structure based on combined MODELLER DOPE and PROCHECK G-FACTOR scores are considered here. (A) Distribution mean = 1.55Å, standard deviation = 0.92Å. (B) Distribution mean = 1.17Å, standard deviation = 0.67Å.

