## Supplementary material for "A Graph-Directed Approach for Creation of a Homology Modeling Library: Application to Venom Structure Prediction": Template Libraries: 6C_rost_table.tex

\begin{table}[]
\centering
\caption{}
\begin{tabular}[t]{p{0.3\linewidth}>{\raggedright\arraybackslash}p{0.3\linewidth}>{\raggedright\arraybackslash}p{0.3\linewidth}}
\toprule
\textbf{Name} & \textbf{PDB ID} & \textbf{Sequence} \\
\midrule
c\-onot\-oxin\--GS & 1\-ag7 & A\-CSGR\-GSRC\-PPQC\-CMGL\-RCGR\-GNPQ\-KCIG\-AHED\-V \\
P\-IIIE\-, PI\-IIE \-[K9S\-], P\-IIIE\- [S1\-7Y,S\-18N,\-S20L\-] & 1\-jlo & H\-PPCC\-LYGK\-CRRY\-PGCS\-SASC\-CQR \\
M\-VIIC\-, S6\-.6 & 1\-omn & C\-KGKG\-APCR\-KTMY\-DCCS\-GSCG\-RRGK\-C \\
T\-VIIA\- & 1\-eyo & S\-CSGR\-DSRC\-PPVC\-CMGL\-MCSR\-GKCV\-SIYG\-E \\
T\-xVII\- & 1\-f3k & C\-KQAD\-EPCD\-VFSL\-DCCT\-GICL\-GVCM\-W \\
T\-xVIA\- & 1\-fu3 & W\-CKQS\-GEMC\-NLLD\-QNCC\-DGYC\-IVLV\-CT \\
E\-VIA & 1\-g1z & D\-DCIK\-PYGF\-CSLP\-ILKN\-GLCC\-SGAC\-VGVC\-ADL \\
G\-IIIB\- & 1\-gib & R\-DCCT\-PPRK\-CKDR\-RCKP\-MKCC\-A \\
G\-VIA & 1\-ttl & C\-KSPG\-SSCS\-PTSY\-NCCR\-SCNP\-YTKR\-CY \\
P\-IVA & 1\-p1p & G\-CCGS\-YPNA\-ACHP\-CSCK\-DRPS\-YCGQ\- \\
E\-IVA & 1\-pqr & G\-CCGP\-YPNA\-ACHP\-CGCK\-VGRP\-PYCD\-RPSG\-G \\
P\-IIIA\- & 1\-r9i & Q\-RLCC\-GFPK\-SCRS\-RQCK\-PHRC\-C \\
M\-VIIA\-[R10\-K] & 1\-tt3 & C\-KGKG\-AKCS\-KLMY\-DCCT\-GSCR\-SGKC\- \\
A\-m276\-6 & 1\-yz2 & C\-KQAG\-ESCD\-IFSQ\-NCCV\-GTCA\-FICI\-E \\
M\-rIII\-E & 2\-efz & V\-CCPF\-GGCH\-ELCY\-CCD \\
F\-VIA & 2\-km9 & C\-KGTG\-KSCS\-RIAY\-NCCT\-GSCR\-SGKC\- \\
I\-m23a\-, Mr\-23a & 2\-lmz & I\-PYCG\-QTGA\-ECYS\-WCIK\-QDLS\-KDWC\-CDFV\-KDIR\-MNPP\-ADKC\-P \\
B\-uIII\-B & 2\-lo9 & V\-GERC\-CKNG\-KRGC\-GRWC\-RDHS\-RCC \\
K\-IIIA\-, KI\-IA [\-W8dT\-rp] & 2\-lxg & C\-CNCS\-SKWC\-RDHS\-RCC \\
A\-r144\-6 & 2\-m61 & C\-CRLA\-CGLG\-CHPC\-C \\
 & 2\-mso & S\-CNNS\-CQSH\-SDCA\-SHCI\-CTFR\-GCGA\-VNGL\-P \\
 & 2\-msq & S\-CGGS\-CFGG\-CWPG\-CSCY\-ARTC\-FRDG\-LP \\
 & 2\-mw7 & D\-GECG\-DKDE\-PCCG\-RPDG\-AKVC\-NDPW\-VCIL\-TSSR\-CENP\- \\
M\-fVIA\- & 2\-n7f & R\-DCQE\-KWEY\-CIVP\-ILGF\-VYCC\-PGLI\-CGPF\-VCV \\
 & 2\-n8e & C\-RIPN\-QKCF\-QHLD\-DCCS\-RKCN\-RFNK\-CVLP\-ETGG\-G \\
 & 2\-n8h & G\-WCGD\-PGAT\-CGKL\-RLYC\-CSGF\-CDSY\-TKTC\-KDKS\-SA \\
C\-nIII\-C & 2\-yen & Q\-GCCN\-GPKG\-CSSK\-WCRD\-HARC\-C \\
C\-cTx & 4\-b1qP\- & A\-PWLV\-PSQI\-TTCC\-GYNP\-GTMC\-PSCM\-CTNT\-C \\
R\-eg12\-i & 6\-bx9 & C\-CTAL\-CSRY\-HCLP\-CC \\
 & 6\-ceg & C\-KPPG\-SKCS\-PSMR\-DCCT\-TCIS\-YTKR\-CRKY\-Y \\
\bottomrule
\end{tabular}
\label{}
\end{table}
